# The long noncoding RNA *Meg3* regulates myoblast plasticity and muscle regeneration through epithelial-mesenchymal transition

**DOI:** 10.1101/2020.06.15.152884

**Authors:** Tiffany L. Dill, Alina Carroll, Jiachen Gao, Francisco J. Naya

## Abstract

Formation of skeletal muscle is among the most striking examples of cellular plasticity in animal tissue development, where mononucleated, lineage-restricted progenitor cells are reprogrammed by epithelial-mesenchymal transition (EMT) to produce multinucleated myofibers. While some mediators of EMT have been shown to function in muscle formation, the regulation of this process in this tissue remains poorly understood. The long noncoding RNA (lncRNA) *Meg3* is processed from the >200 kb *Dlk1-Dio3* polycistron that we have previously shown is involved in skeletal muscle differentiation and regeneration. Here, we demonstrate that *Meg3* regulates EMT in myoblast differentiation and skeletal muscle regeneration. Chronic inhibition of *Meg3* in C2C12 myoblasts promoted aberrant EMT activation, and suppressed cell state transitions required for fusion and myogenic differentiation. Furthermore, adenoviral *Meg3* knockdown compromised muscle regeneration, which was accompanied by abnormal mesenchymal gene expression and interstitial cell proliferation in the regenerating milieu. Transcriptomic and pathway analyses of *Meg3*-depleted C2C12 myoblasts and injured skeletal muscle revealed a significant dysregulation of EMT-related genes, and identified TGFβ as a key upstream regulator. Importantly, chemical inhibition of TGFβR1, as well as its downstream effectors ROCK1/2 and p38 MAPK, restored many aspects of myogenic differentiation in *Meg3*-depleted myoblasts *in vitro*. Thus, *Meg3* regulates myoblast identity to maintain proper cell state for progression into differentiation.

**Summary statement:** Muscle differentiation and regeneration are regulated by an evolutionarily conserved long noncoding RNA that restricts gene expression to coordinate cell state transitions

## Introduction

During developmental myogenesis, polarized epithelial cells within mesoderm-derived somites undergo dramatic changes in cell identity through epithelial-mesenchymal transition (EMT) to generate muscle progenitors with the ability to migrate and ultimately fuse to form multinucleated myotubes (Yusuf and Brand-Saberi 2006; Buckingham and Rigby 2014; Chai and Pourquie 2017). Similarly, repair of damaged or diseased skeletal muscle requires extensive cell state transitions in muscle and non-muscle progenitors, which collaborate to form newly functional myofibers (Uezumi et al 2014; Wosczyna and Rando 2018). The temporal control of EMT in both embryonic and regenerative myogenesis is essential for proper muscle formation, and while numerous effectors of EMT participate in myogenic differentiation (Kollias and McDermott 2008; Krauss 2010), less is known about regulatory factors that coordinate the various facets of this process.

The imprinted *Dlk1-Dio3* locus is the largest known mammalian cluster of noncoding RNAs (ncRNAs), encoding three annotated lncRNAs (*Meg3*, *Rian*, *Mirg*), numerous small nucleolar RNAs (snoRNAs), and over 60 miRNAs co-transcribed from the maternal allele (da Rocha *et al.* 2008; Benetatos *et al.* 2013; Dill and Naya 2018). In recent years, this locus has emerged as a key regulator of cellular processes as diverse as pluripotency and metabolism (Stadtfeld *et al.* 2010; Liu *et al.* 2010; Benetatos *et al.* 2014; Kameswaran *et al.* 2014; Qian *et al.* 2016). In muscle, *Dlk1-Dio3* miRNAs function downstream of the MEF2A transcription factor to modulate WNT signaling in differentiation and regeneration, and to regulate mitochondrial activity in satellite cells (Snyder *et al.* 2013; Wust *et al.* 2018; Castel *et al.* 2018). Dysregulation of *Dlk1-Dio3* ncRNA expression has also been documented in muscular dystrophies, sarcopenia, and in rare congenital growth disorders with pleiotropic organ defects including hypotonia (Eisenberg *et al.* 2007; Ogata *et al.* 2008; Ioannides *et al.* 2014; Mikovic *et al.* 2018). Interestingly, a partial deletion of the *Dlk1-Dio3* locus at the 3’end – harboring 22 miRNAs (miR-379/miR-544) – in mice resulted in skeletal muscle hypertrophy (Gao *et al.* 2015), but not in mice with an additional 17 miRNAs removed from the 3’ end (miR-379/miR-410) (Labialle *et al.* 2014). Given the vast array of ncRNAs with distinct regulatory activities encoded by this massive and unusually complex locus, much remains to be learned about the individual versus collective function of these ncRNAs.

Recently, the Tabula Muris Consortium performed a large-scale single-cell RNA sequencing (scRNA-seq) screen of numerous postnatal mouse tissues including skeletal muscle (Schaum *et al.* 2018). Although the syncytial nature of skeletal myofibers precluded their analysis by scRNA-seq, this procedure effectively screened multiple mononucleated cells known to be present in the interstitium of this tissue. Analysis of the single-cell muscle transcriptome from this database revealed significantly enriched transcripts of the *Dlk1-Dio3* encoded lncRNA *Meg3* (also known as *Gtl2*) in satellite cells and mesenchymal stromal cells; these transcripts were largely absent from fibroblasts, endothelial cells and cells of the immune system including macrophages, B-cells, and T-cells (Schaum *et al.* 2018). To date, *Meg3* is one of the most comprehensively characterized *Dlk1-Dio3* ncRNAs and has been shown to interact with *Polycomb* repressive complex 2 (PRC2) to regulate chromatin structure in pluripotent embryonic stem cells and neuronal cells (Zhao *et al.* 2010; Kaneko *et al.* 2014; Das *et al.* 2015; Yen Y-P *et al.* 2018). An epigenetic function of *Meg3* has also been reported in human breast and lung cancer cells, where it suppresses TGFβ-related genes to modulate their invasive characteristics (Mondal *et al.* 2015; Terashima *et al.* 2017).

Given the enrichment of *Meg3* in muscle satellite cells and its epigenetic function, we hypothesized that *Meg3* regulates cell identity in myoblasts during differentiation and regeneration. To this end, we performed *Meg3* knockdown in C2C12 myoblasts and regenerating skeletal muscle to examine for changes in cell identity and mesenchymal character. For both model systems, *Meg3* inhibition upregulated EMT-related genes resulting in enhanced mesenchymal characteristics and impaired myotube formation, illuminating a key role for *Meg3* muscle cell identity *in vitro* and *in vivo*.

## Results

### Chronic *Meg3* knockdown in myoblasts impairs myotube formation

To investigate the role of *Meg3* in regulating myoblast identity, we first examined its expression in various muscle-derived cells. Single-cell RNA sequencing data by *Tabula muris* (Schaum *et al.* 2018) revealed that *Meg3* expression was restricted to satellite and mesenchymal cells in adult mouse limb muscle, and more enriched than the other two annotated *Dlk1-Dio3* locus lncRNAs, *Rian* and *Mir*g (Fig. 1A). Moreover, *Meg3* was highly upregulated in injury-induced muscle regeneration (Fig. 1B), and enriched in proliferating C2C12 myoblasts, but downregulated upon differentiation (Fig. 1C). Taken together, these data suggest *Meg3* functions in muscle and non-muscle progenitor cells.

**Figure 1.**
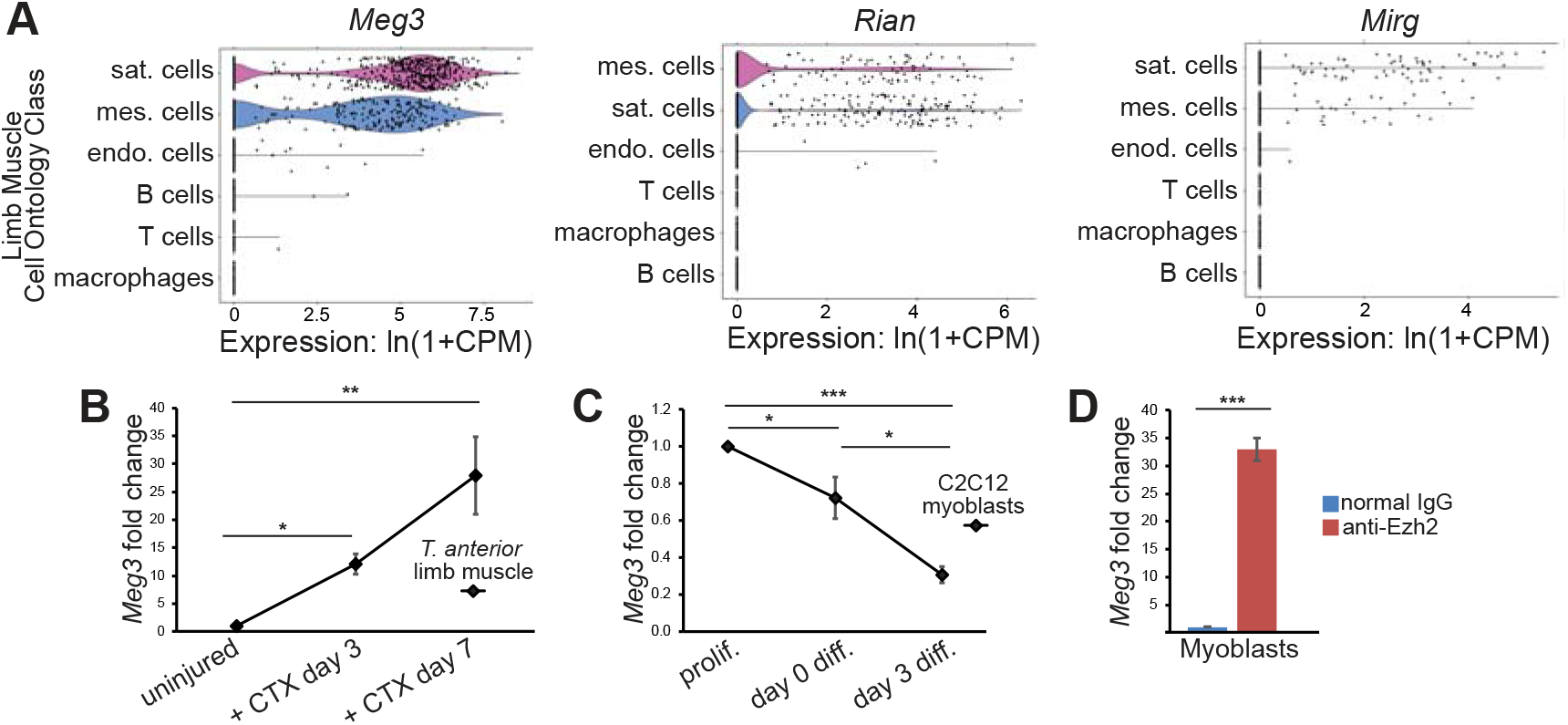
*Meg3* lncRNA is enriched in muscle progenitors and mesenchymal stromal cells. **A)** Violin plots depict limb muscle single cell RNAseq data for *Dlk1*-*Dio3* megacluster-encoded lncRNAs *Meg3*, *Rian*, and *Mirg*. All lncRNAs showed enrichment in satellite (sat.) and mesenchymal (mes.) cell types, with *Meg3* as the most abundant. CPM = counts per million reads mapped. **B)** qPCR temporal *Meg3* expression profiling was performed on regenerating mouse *Tibialis anterior* (TA) muscle tissue harvested before (uninjured), and on the indicated days after cardiotoxin injection (+CTX). *Meg3* lncRNA transcripts were upregulated following CTX-induced injury, which corresponds with satellite and mesenchymal cell expansion (n=3 mice per time point). **C)** qPCR temporal expression profiling of *Meg3* in C2C2 myoblast differentiation. *Meg3* transcripts were most enriched during proliferation (prolif.), and progressively downregulated during course of differentiation (n=4). **D)** RNA immunoprecipitation (RNA-IP) was performed on subconfluent C2C12 myoblast lysates to examine for *Meg3*-PRC2 interaction. Immunoprecipitated RNA was quantified by qPCR, using supernatant as an internal normalization control. Compared to normal IgG controls, *Meg3* was enriched in anti-Ezh2 immunoprecipitates (n=3 sets of 60 plates).

Based on the established interaction between *Meg3* and the PRC2 subunit Ezh2 (Zhao *et al.* 2010; Kaneko *et al.* 2014; Das *et al.* 2015), we performed native RNA-immunoprecipitation (RNA-IP) to determine whether *Meg3* interacts with this *Polycomb* complex in proliferating myoblasts. *Meg3* transcripts were enriched 30-fold in Ezh2 immunoprecipitates from C2C12 myoblasts (Fig. 1D), demonstrating this epigenetic interaction is conserved in muscle progenitor cells.

We next inhibited *Meg3* in C2C12 myoblasts to determine whether *Meg3* is required for muscle differentiation. Given that the *Dlk1-Dio3* ncRNAs are transcribed as a large polycistronic primary RNA (Tierling et al 2006; Zhou et al 2010; Luo et al 2016), processed *Meg3* transcripts were directly targeted with *Meg3*-specific short hairpin RNA (shRNA) (Mondal et al 2015) to circumvent unwanted effects on expression of adjoining RNAs in the cluster. Transfected C2C12 myoblasts were subjected to G418 selection, and clonal populations were isolated for differentiation experiments. Both clonal and non-clonal (heterogeneous) populations of myoblasts with stably integrated sh*Meg3* displayed significantly reduced *Meg3* expression compared to sh*LacZ* control myoblasts (Fig. 2A).

**Figure 2.**
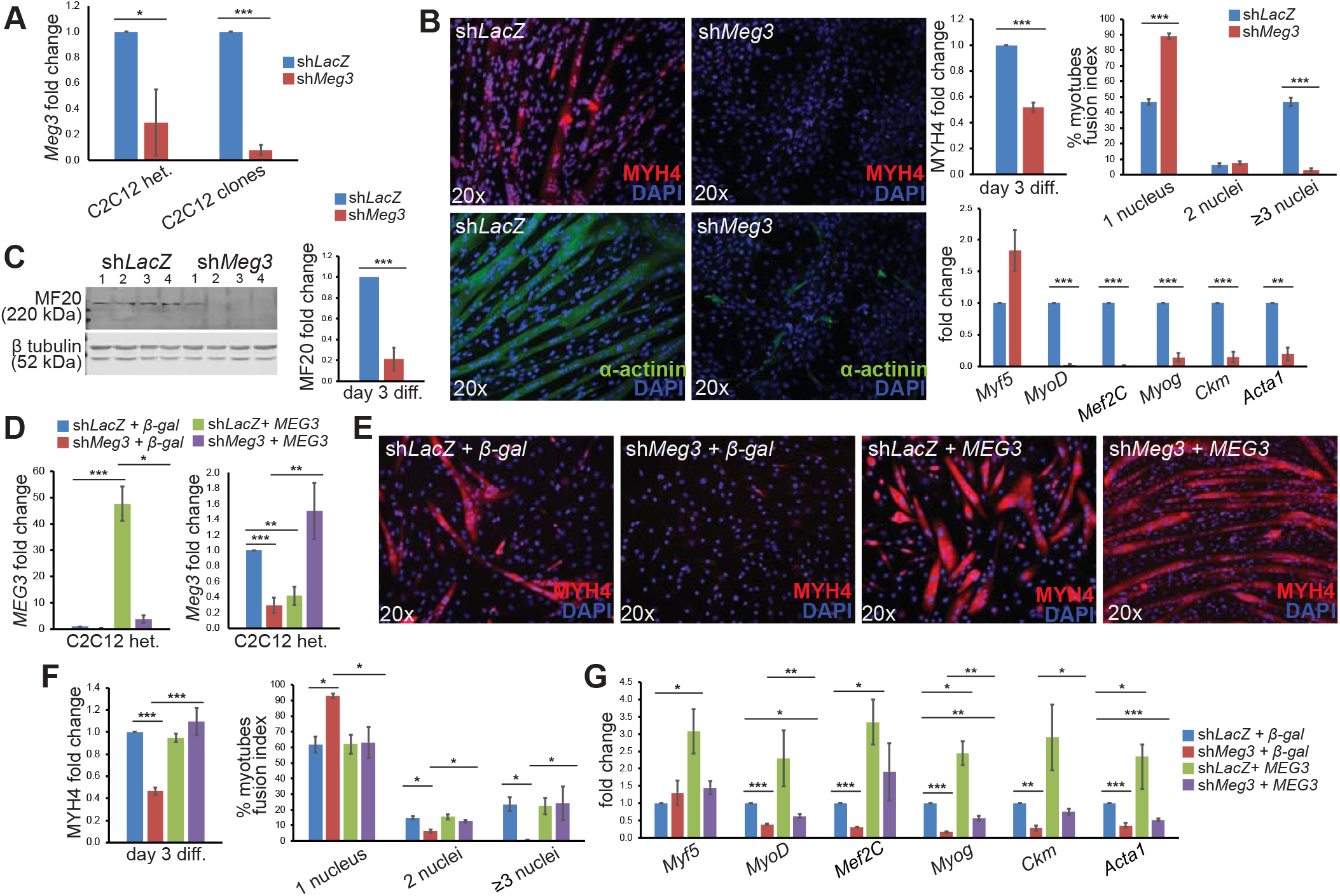
*Meg3* is required for C2C12 myoblast differentiation. **A)** qPCR quantification of *Meg3* transcript levels in heterogeneous cell populations derived from G418 selection (C2C12 het.), and subsequently derived clonal populations (C2C12 clones) indicate that stable shRNA integration resulted in *Meg3* knockdown (n=3). **B)** Immunofluorescence quantification of MYH4 indicates markedly reduced expression in sh*Meg3* C2C12 clones. Quantification of nuclei within α-actinin cell-boundaries show reduced fusion index in sh*Meg3* clones (n=3). qPCR expression profiling indicated unchanged *Myf5* transcript levels, but significant reduction in other myogenic differentiation markers (n=3). **C)** Western blot quantification of MF20 signal (normalized to β-tubulin) showed marked reduction specific to sh*Meg3* clones (n=4). **D)** qPCR quantification confirmed overexpression of human *MEG3* in sh*LacZ* and sh*Meg3* myoblasts, and restoration of endogenous *Meg3* transcript levels relative to β-galactosidease controls (n=3). **E)** Human *MEG3* restored both MYH4 expression and fusion index in sh*Meg3* but not sh*LacZ* myoblasts (n=6 MYH4, n=3 fusion index). **F)** qPCR expression profiling of heterogeneous rescue clones revealed an increase in *Mef2C, Ckm*, *MyoD* and *Myog* levels in sh*Meg3* + *MEG3* myotubes. *MYH4* = myosin heavy chain 4 (Proteintech antibody), MF20 = myosin heavy chain 4 (DHSB antibody), *Myf5* = *myogenic factor 5*, *MyoD* = myogenic differentiation 1, *Mef2C* = *myogenic enhancing factor-*2 *C*, Myog = *Myogenin*, *Ckm* = *Muscle Creatine Kinase*, *Acta1* = skeletal muscle actin.

To determine the functional consequences of chronic *Meg3*-depletion on myoblast differentiation, the ability of C2C12 sh*Meg3* and sh*LacZ* clones to form myotubes was examined. Three days after initiating differentiation sh*LacZ* clones formed numerous multinucleated myotubes with robust expression of myosin heavy chain (MYH4), a differentiation marker expressed in myotubes but not myoblasts (Fig. 2B). In contrast, MYH4-expressing myotubes were largely absent in sh*Meg3* clones, and levels of this protein were dramatically reduced (Fig. 2B and C). Given the difficulty of detecting MYH4 expression in *Meg3*-deficient myoblasts, we immunostained with α-actinin to quantify fusion index, a readout of myotube formation. Fusion was markedly reduced in sh*Meg3* myotubes, with nearly 90% of cells remaining mononucleated, and less than 5% containing greater than three nuclei (Fig. 2B, upper graph). In stark contrast, nearly half of all sh*LacZ* control myotubes contained three or more nuclei. Notably, extending the differentiation timeline to day 7 did not improve myotube formation (data not shown).

Supporting the above observations, transcripts for the myogenic regulatory factors *MyoD*, *myogenin*, and *Mef2c*, and terminal differentiation marker genes *muscle creatine kinase* and *skeletal α-actin* were significantly reduced in sh*Meg3* myotubes (Fig. 2B, lower graph). These results reinforce the observation that *Meg3* deficiency severely impairs myogenic differentiation.

Rescue experiments were subsequently performed to demonstrate the specificity of the *Meg3* knockdown phenotype. Because overexpression of human *MEG3*, either transiently or permanently integrated through a second selection process (puromycin), failed to rescue differentiation in established sh*Meg3* myoblast lines (data not shown), stable C2C12 myoblasts were generated *de novo* by co-transfection of human *MEG3* or β-galactosidase and either sh*Meg3* or sh*LacZ* expression plasmids. Following G418 selection, expression of human *MEG3* was examined in these co-transfections and found to be increased more than 50-fold in sh*LacZ* + *MEG3* controls, but increased only 3-fold in sh*Meg3* + *MEG3* myoblasts (Fig. 2D). Comparatively low *MEG3* levels in myoblasts co-transfected with sh*Meg3* was not unexpected, as the sh*Meg3* hairpin targets an evolutionarily conserved *Meg3* sequence shared by mouse, rat, and human transcripts. Examination of endogenous *Meg3* expression in sh*Meg3* + *MEG3* myoblasts also revealed that these transcripts were restored to relatively normal levels (Fig. 2D). Stably integrated myoblasts were subsequently induced to differentiate, which resulted in rescue of cytosolic MYH4 expression and fusion index in clones harboring co-integration of sh*Meg3* and human *MEG3* (Fig. 2E and F). In agreement with improved myotube formation, expression of *MyoD*, *myogeni*n, *Mef2c*, and *muscle creatine kinase* was augmented in sh*Meg3* myoblasts by overexpression of human *MEG3*, but not to the extent observed in sh*LacZ* controls (Fig. 2G). These results clearly demonstrate that *Meg3* functions in myogenic differentiation, and indicates the sh*Meg3*-induced phenotype is not caused by secondary off-target effects.

### Differentiation defect in *Meg3*-deficient myoblasts is associated with reduced proliferation, survival and mitochondrial activity

*Meg3* is predicted to function as a tumor-suppressor via its interaction with p53 (Zhou *et al.* 2007), leading us to speculate that *Meg3* could regulate proliferation in myoblasts. During expansion of sh*Meg3* and sh*LacZ* clones we noted that *Meg3*-deficient cells grew more slowly than control myoblasts. To determine whether *Meg3* depletion modulates proliferation, we subjected clones to BrdU incorporation assays and found that sh*Meg3* clones exhibit markedly reduced proliferation (Fig. 3A). Moreover, assays for cleaved caspase 3 (Fig. 3B, left) and cell viability (Fig. 3B, right) strongly suggest that *Meg3* depletion also resulted in increased cell death.

**Figure 3.**
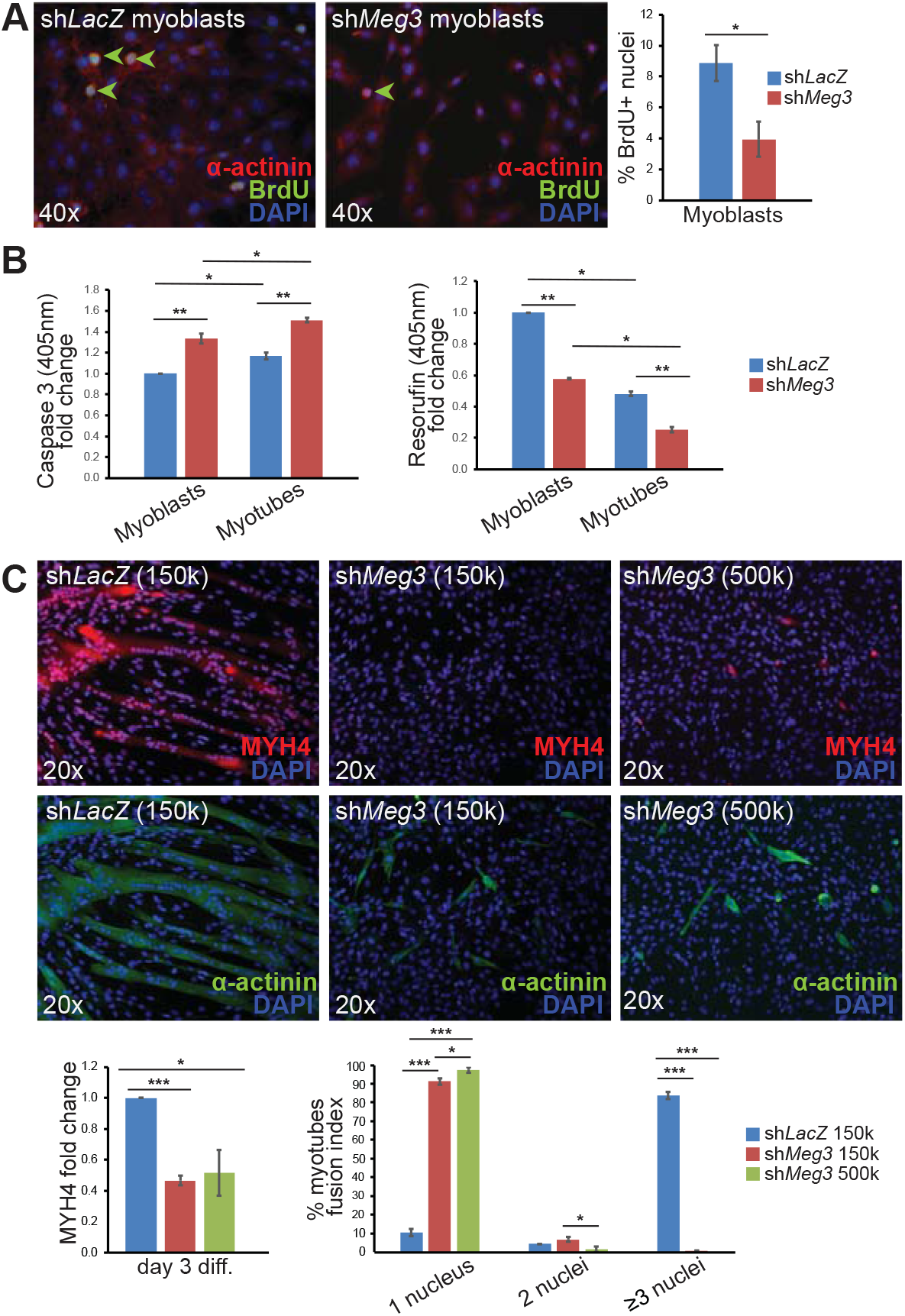
*Meg3* modulates C2C12 myoblast proliferation and viability. **A)** Quantification of BrdU+ nuclei (green arrowheads) indicated that sh*Meg3* myoblasts divided at a reduced frequency (n=3). **B)** Cleaved caspase 3 assay revealed higher apoptosis in sh*Meg3* myoblasts and myotubes relative to control (n=3). Cell Titer Blue viability assay indicated reduced viability in sh*Meg3* myoblasts and myotubes relative to control (n=3). **C)** sh*Meg3* myoblasts seeded at increasing densities was not sufficient to restore MYH4 expression or fusion index (n=3).

Reasoning that impaired myogenic differentiation could be an artifact of low myoblast density, we seeded sh*Meg3* clones at high numbers and subsequently examined for their ability to differentiate. However, impaired myogenic differentiation of sh*Meg3* cells persisted even with increased seeding density, as indicated by low cytoplasmic MYH4 expression and fusion index (Fig. 3C). Taken together, these data suggest that *Meg3* is required for normal cell expansion and viability, but failure to form myotubes in *Meg3* deficient myoblasts is not a consequence of reduced cell-cell contacts.

Because *Dlk1-Dio3* locus expression levels have been shown to correlate with changes in mitochondrial activity in satellite cells (Wust *et al.* 2018) we also examined whether *Meg3* depletion affected mitochondria in C2C12 myoblasts. Mutant and control clones were pulsed with MitoTrackerCMXRos and analyzed for mitochondrial mass as myoblasts or myotubes. While Mitotracker Red CMXRos signal increased in response to differentiation conditions within each treatment group, sh*Meg3* clones displayed lower mitochondrial signal when compared with sh*LacZ* clones for early but not late timepoints (Supplemental Fig. 1). These data suggest that *Meg3* is required for maintaining normal mitochondrial activity in proliferating, but not differentiating myoblasts.

### Epithelial-mesenchymal transition (EMT) is regulated by *Meg3* and its inhibition alters myoblast identity

To understand the molecular mechanisms of differentiation impairment in sh*Meg3* myoblasts in an unbiased manner, RNA-sequencing (RNA-seq) (Illumina) was performed. Stable *Meg3* knockdown had a profound effect on gene expression, resulting in thousands of transcripts differentially expressed on differentiation day 3 (Fig. 4A). Despite the expected repressive function of *Meg3* based on its interaction with Ezh2 in C2C12 myoblasts, a similar number of up- and down-regulated transcripts were observed (Fig. 4A). This result is consistent with a satellite cell-specific knockout of Ezh2, which did not result in global derepression of the transcriptome (Juan *et al.* 2011). The near complete inability to form multinucleated myotubes is reminiscent of defects in myoblast fusion pathways, yet expression profiling of master regulators of myoblast fusion such as *myomaker* and *myomixer* were not significantly downregulated (Supplemental Fig. 2). Curiously, a few fusion genes such as *Kirrel*, *Ehd2*, and *Fer1L5*, were found to be upregulated in *Meg3*-deficient myoblasts suggesting that any effect on fusion would be enhanced rather than impaired.

**Figure 4.**
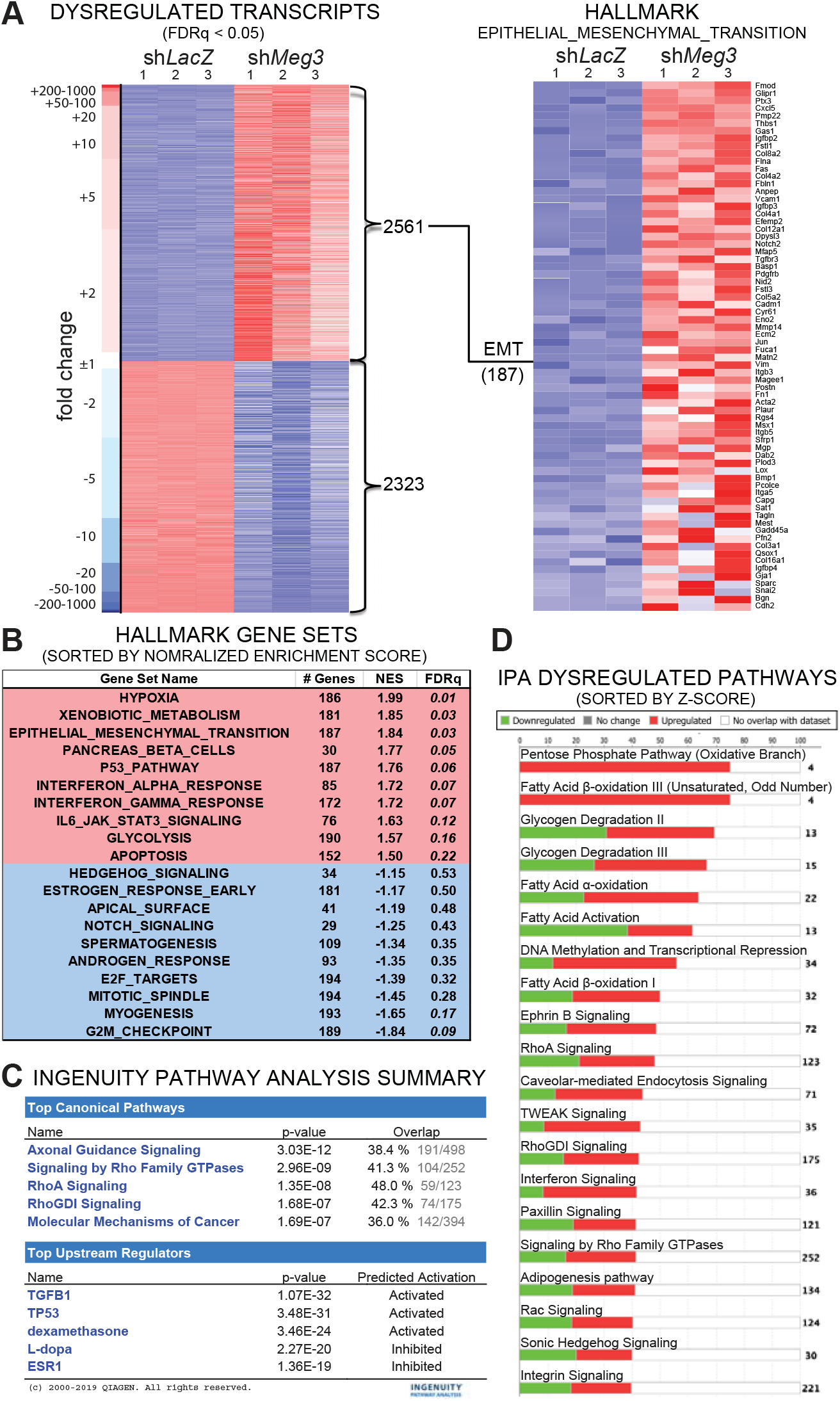
Chronic *Meg3* knockdown in C2C12 myoblasts is associated with activation of EMT and TGFβ. **A)** Heatmap of 4884 dysregulated transcripts, with 2561upregulated (red) and 2323 downregulated (blue). Broad GSEA Hallmark analysis indicated coherent upregulation of 187 well-characterized EMT gene markers. **B)** Table conveying the top 10 up (red) and down (blue) - regulated Hallmark biological states and cellular processes, as indicated by normalized enrichment score (NES). EMT is among the top three upregulated gene sets. **C)** Summary output from Qiagen Ingenuity Pathway Analysis (IPA) software performed with significantly dysregulated transcripts with p<0.05. IPA lists TGFβ1 activation as a Top Upstream Regulator, and lists various Rho family pathways among the Top Canonical Pathways. **D)** Stacked bar graphs indicate proportions of upregulated (red) and downregulated (green) genes that comprise the top 20 dysregulated biological pathways, which include RhoA-related pathways (RhoA signaling, RhoGDI signaling, Signaling by Rho Family GTPases, Rac signaling).

Gene set enrichment analysis (GSEA) of upregulated transcripts in *Meg3*-deficient myoblasts revealed significant enrichment in metabolism (glycolytic), epithelial-mesenchymal transition (EMT), p53 signaling, and apoptosis (Fig. 4B). In contrast, downregulated transcripts were enriched for cell cycle, myogenesis, Notch signaling, and apical surface (Fig. 4B). While the downregulation of myogenesis was entirely consistent with the phenotype of *Meg3*-depleted myoblasts, the GSEA results highlight a broader gene regulatory effect of *Meg3* on cellular processes. To obtain a more focused picture of altered pathways in *Meg3*-deficient myoblasts, we also performed Ingenuity Pathway Analysis (IPA). This analysis revealed a significant enrichment in axonal guidance and Rho signaling, and predicted TGFβ1 as the top upstream regulator of gene expression changes (Fig. 4C and D). Based on the relationship between TGFβ and Rho activity in cytoskeletal remodeling in cell migration (Ungefroren *et al.* 2018), the integrated GSEA and IPA results indicated that EMT is a key pathway affected by *Meg3*-deficiency in C2C12 myoblasts.

We next verified expression levels of genes belonging to EMT, primarily focusing on cellular processes integral to this pathway such as cell adhesion, migration, and cytoskeletal remodeling. Initially, expression of cadherins, key mediators of calcium-dependent cell adhesion, was examined for cadherin switching – a well-characterized signature of EMT where N-cadherin becomes enriched (Lamouille *et al.* 2014). As shown in Fig. 5A, expression of mesenchymal *N-cadherin* (*Cdh2*) transcript and protein levels were significantly upregulated in *Meg3* knockdown cells. Although we observed significant downregulation of epithelial *E-cadherin* (*Cdh1*) transcripts, protein was difficult to detect by western blot analysis given the lower sensitivity of this assay and mesenchymal character of myoblasts. These data suggest that *Meg3* knockdown myoblasts have undergone cadherin-switching.

**Figure 5.**
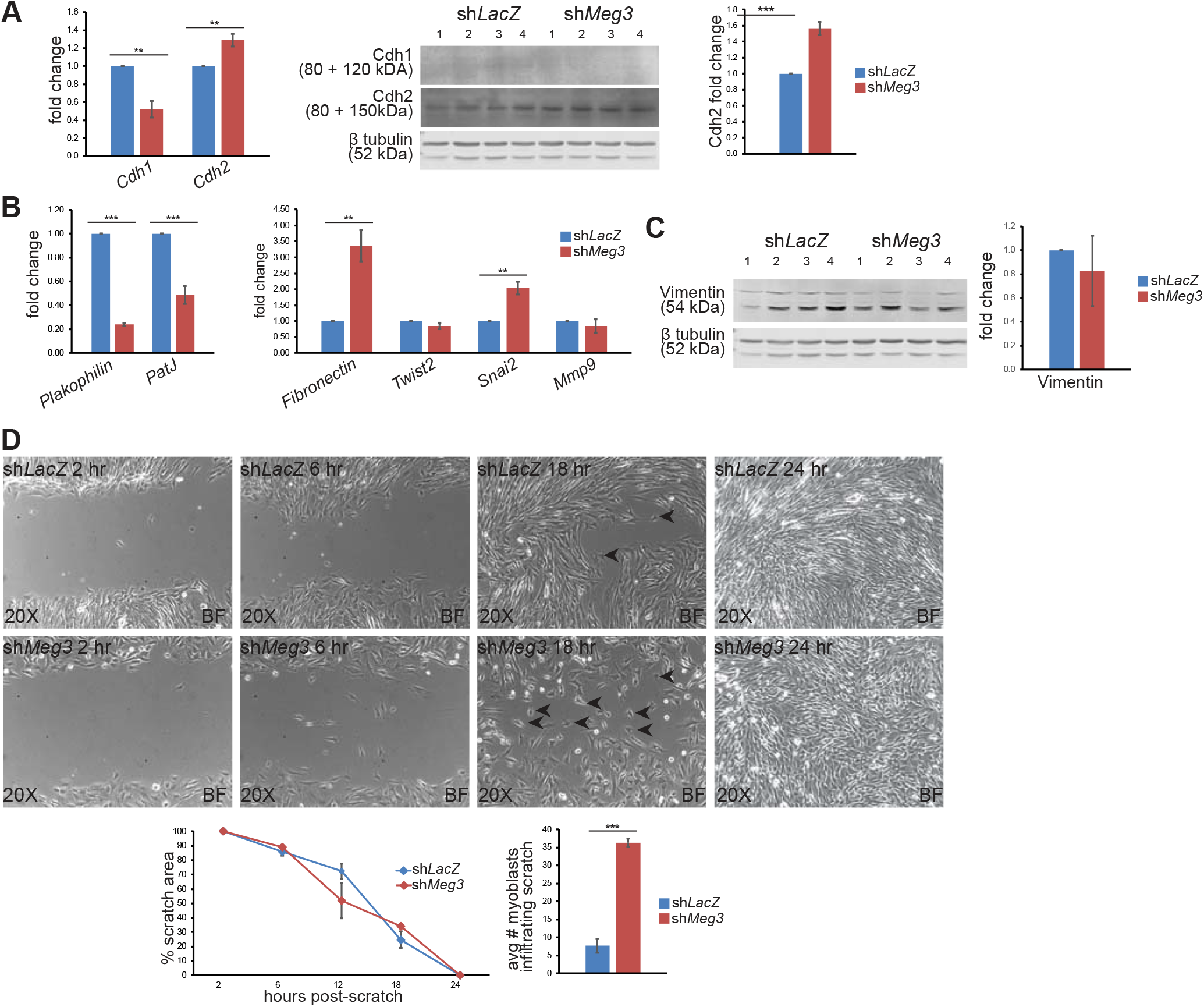
*Meg3* knockdown enhances the mesenchymal character of C2C12 myoblasts. **A)** Cadherin switching was assessed by qPCR (left) and Western blot (right) quantification of E-cadherin (Cdh1) and N-cadherin (Cdh2) in day 3 myotubes. *Cdh1* transcripts were significantly downregulated in sh*Meg3* myotubes relative to sh*LacZ* controls (n=3), but Cdh1 protein signal from myotubes lysates was extremely faint (n=4). *Cdh2* transcripts were upregulated (n=3), as was Cdh2 protein (n=4). **B)** qPCR expression profiling indicated significant downregulation of epithelial *Plakophilin* and *PatJ* transcripts in sh*Meg3* myotubes, with simultaneous upregulation of mesenchymal *Fibronectin* and *Snai2* (n=3). **C)** Western blot of Vimentin normalized to β-tubulin indicated no change in mesenchymal Vimentin (n=4). **D)** Scratch-wound assays revealed no detectable change in wound-healing efficiency, as measured by % scratch area (n=3). Brightfield (BF) microscopy of wound-healing morphology differed between sh*LacZ* control and sh*Meg3* clones, with a higher proportion of myoblasts invading the scratch territory with fewer than 2 cell contacts (black arrowheads, n=3).

Subsequently, expression profiling of selected epithelial and mesenchymal marker genes was examined. Both *Plakophilin* (a cytoskeletal protein that anchors cadherins to intermediate filaments) and *PatJ* (a cell-polarity protein crucial for tight-junctions) were downregulated in *Meg*3 knockdown cells, while mesenchymal markers *Fibronectin* (an integrin-binding extracellular matrix glycoprotein) and *Snail2* (a transcriptional repressor of epithelial genes in developmental cell migration and cancer) were both upregulated (Fig. 5B). *Meg3* knockdown did not affect transcript levels of *Twist2* (a bHLH transcription factor associated with carcinogenic EMT) or *Mmp9* (an adhesion-related matrix metallopeptidase) (Fig. 5B), or had a significant effect on expression of vimentin (an intermediate filament protein that modulates cell shape and motility) (Fig. 5C). Taken together, the bioinformatics and molecular analyses of the sh*Meg3* transcriptome reveal reprogramming of epithelial and mesenchymal gene expression patterns, which strongly suggests activation of EMT and an identity shift towards enhanced mesenchymal character.

### *Meg3*-deficient myoblasts display mesenchymal-like cell adhesion and migration properties

We next sought to investigate cell behavior associated with EMT such as adhesion and migration in greater detail. To analyze migration, a property enhanced in mesenchymal cells, we performed scratch wound assays to determine whether the observed perturbations in EMT affected this cellular behavior. Measurement of wound closure using the Volkner Wound Healing Tool (ImageJ) indicated no difference in the rate of wound-closure between control and *Meg*3 knockdown C2C12 cells (Fig. 5D). However, the number of infiltrating cells sharing fewer than two borders with neighboring cells was significantly higher in sh*Meg3*-myoblasts, which was most pronounced at the 18hr timepoint (Fig 5D). By contrast, sh*LacZ* control myoblasts formed sheets reminiscent of collective migration behavior (Campbell and Casanova 2016). Curiously, we also noted that sh*Meg3* myoblasts required more time to trypsinize during passaging, which may reflect perturbed adhesion properties – and plating these cells on a glass substrate slightly improved fusion of cells with 2 nuclei, but not ≥3 nuclei (Supplemental Data Fig 3). These observed changes in migration and adhesion characteristics indicate that activation of EMT in sh*Meg3* myoblasts facilitated mesenchymal behaviors that deviate from normal myoblasts.

### Inhibition of the TGFβ signaling pathway restores fusion competence to sh*Meg3* mutant myoblasts

Considering TGFβ1 emerged as a top upstream regulator (IPA) and its activity is known to inhibit myogenesis (Kollias and McDermott 2008), we investigated whether chemical inhibition of this pathway impacts sh*Meg3* myoblast differentiation. Chemical inhibition of TGFβR1 with LY-2157299 (LY) was sufficient to increase C2C12 sh*Meg3* fusion (cells with 3 or more nuclei), and increased MYH4 immunofluorescence signal in α-actinin positive cells (Fig. 6A and B). While these differentiation features were greatly improved compared to sh*Meg3* myoblasts, the fusion index was still lower than sh*LacZ* controls, indicating that fusion competence was only partially restored. Nevertheless, myosin expression (MF-20) was restored in LY-treated sh*Meg3* myoblasts (Fig. 6B), and myogenic transcripts *Mef2C, Myog*, *Ckm*, and *Acta1* were enhanced in sh*Meg3* myotubes (Fig. 6C). Interestingly, LY treatment resulted in upregulation of both epithelial and mesenchymal EMT markers in sh*LacZ* and sh*Meg3* myotubes, but the extent of dysregulation differed (Supplemental Fig. 4). In general, sh*Meg3* cells displayed less induction of epithelial, and greater activation of mesenchymal genes. While LY treatment downregulated vimentin in sh*LacZ* cells, this reduction did not occur in sh*Meg3* cells, suggesting a bias towards retention of mesenchymal character (and more permissive to mesenchymal gene expression) that was independent of rescuing myotube formation.

**Figure 6.**
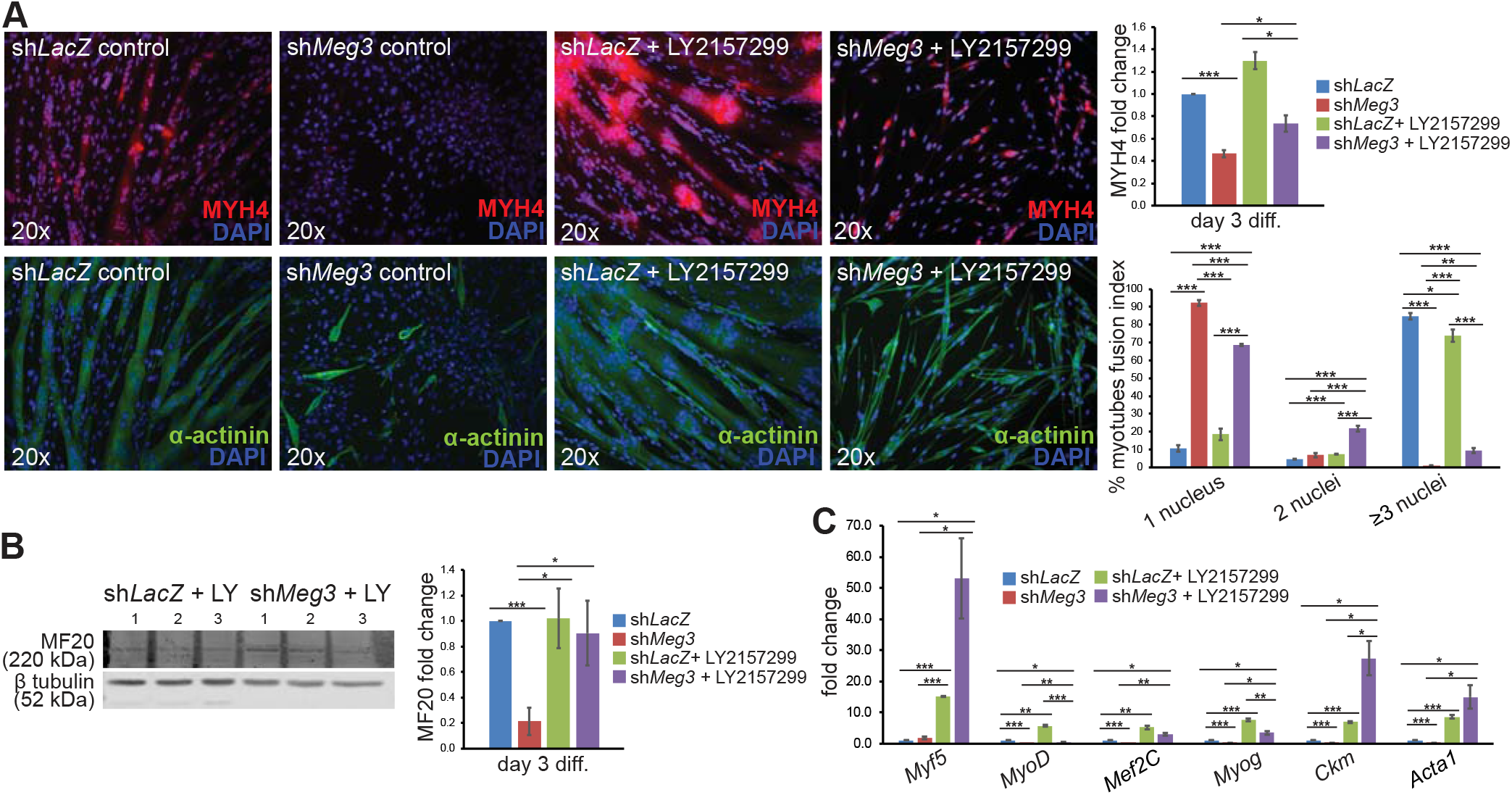
Pharmacological inhibition of TGFβR1 restores fusion competence and myogenic differentiation in *Meg3*-deficient myoblasts. **A)** Following incubation with 10μM LY2157299 (LY), myoblasts were induced to differentiate and examined for MYH4 expression and fusion index. LY-treated sh*Meg3* myotubes adopted an elongated morphology, and displayed increased MYH4, decreased myotubes with 1-nuclei, and increased myotubes with 2- and ≥3 nuclei relative to untreated sh*Meg3* controls (n=3). **B)** Western blot for MF20 indicated that LY-treatment restored myosin heavy chain 4 expression to sh*Meg3* myotubes (n=3). **C)** qPCR expression profiling of LY-treated myotubes indicate significant upregulation of all myogenic markers surveyed (*Myf5*, *MyoD*, *Mef2C*, *Myog*, *Ckm*, *Acta1*) relative to untreated sh*LacZ* myotubes.

The GTPase RhoA functions downstream of TGFβ to remodel the actin cytoskeleton in EMT (Ungefroren *et al.* 2018), and IPA indicated this non-canonical TGFβ signaling effector was dysregulated in sh*Meg3* C2C12 cells (Fig. 4C, D). Considering Rho signaling has been shown to regulate myotube formation (Nishiyama *et al.* 2004) we treated *Meg3*-deficient myoblasts with ROCK1/2 inhibitor Y-27632 (Y), a RhoA-dependent protein kinase, and examined MYH4 expression and fusion index. Consistent with the inhibition of TGFβR1, Y-27632 treatment improved myoblast fusion, but did not improve MYH4 expression in sh*Meg3* cells (Fig. 7A).

**Figure 7.**
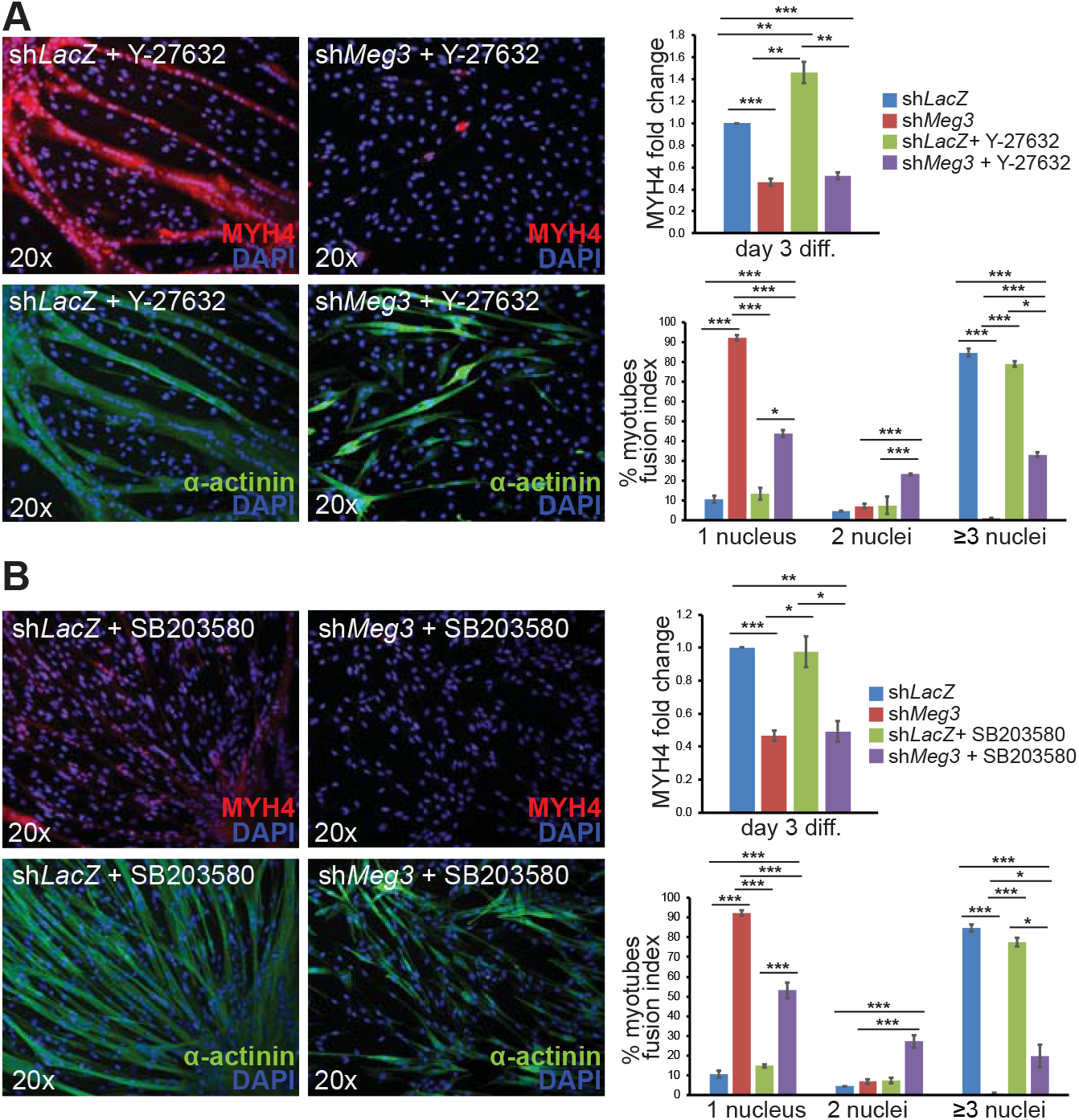
Pharmacological inhibition of Rho GTPase and p38 MAPK restores myotube formation. **A)** Following incubation with 40μM ROCK1/2 inhibitor (Y-27632), myoblasts were induced to differentiate and examined for MYH4 expression and fusion index. Y-27632-treated sh*Meg3* myotubes adopted an elongated morphology, and fusion quantification indicated decreased myotubes with 1-nuclei, and increased myotubes with 2- and ≥3 nuclei relative to sh*Meg3* control (n=3). While MYH4 expression was enhanced in sh*LacZ* + Y-27631 myotubes, MYH4 levels remained unchanged with Y-27632 treatment in sh*Meg3* cells (n=3). **B)** Following incubation with 5μM p38 inhibitor (SB203580), myoblasts were induced to differentiate and examined for MYH4 expression and fusion index. SB203580-treated sh*Meg3* myotubes adopted an elongated spindle-like morphology, and fusion quantification indicated decreased myotubes with 1-nuclei, and increased myotubes with 2- and ≥3 nuclei relative to sh*Meg3* control (n=3). MYH4 expression was unaffected by SB203580 treatment (n=3).

An additional non-canonical TGFβ effector, p38MAPK, has established functions in promoting EMT and modulating myogenic differentiation of satellite cells (Segales *et al.* 2016, Lamouille *et al.* 2014). We treated sh*Meg3* myoblasts with SB-203580, a specific p38 inhibitor, which resulted in improved fusion index, but similar to ROCK1/2 inhibition, did not improve MYH4 expression (Fig. 7B). Taken together, these data strongly suggest that RhoA and p38MAPK signaling modulate fusion in myoblasts, and that other effectors downstream of TGFβ likely regulate MYH4 expression.

While TGFβ is a negative regulator of muscle growth, BMP signaling is thought to work antagonistically to TGFβ, and has been shown to have pro-myogenic effects including muscle mass enhancement (Sartori *et al.* 2013). However, we found that sh*Meg3* myoblasts were largely unresponsive to BMP4 treatment, which was not sufficient to improve MYH4 expression or fusion (Supplemental Fig. 4). These findings indicate that TGFβ-mediated suppression of myogenic differentiation likely overrides BMP4-mediated myogenic enhancement, and reinforces that abnormal TGFβ activation is the primary bottleneck for sh*Meg3* myogenic differentiation.

### *Meg3* knockdown in injured skeletal muscle severely disrupts the normal regenerative response

Based on the enrichment of *Meg3* transcripts in satellite cells and mesenchymal stromal cells we reasoned that this lncRNA functions in the injured skeletal muscle microenvironment, and regulates the identity of these progenitor cell types for proper myofiber regeneration. To circumvent the perinatal lethality of *Meg3* knockout mice (Zhou *et al.* 2010) and other aforementioned effects relating to the polycistron, we used shRNA to knock down *Meg3* in adult skeletal muscle. To this end, *Tibialis anterior* (TA) muscles of mice were injured by cardiotoxin (CTX) co-injected with either sh*Meg3*- or sh*LacZ*-specific adenovirus. Regenerating TA muscles were harvested for expression analysis, and significant downregulation of *Meg3* was confirmed in muscle at 3 days post-injury (Fig. 8A).

**Figure 8.**
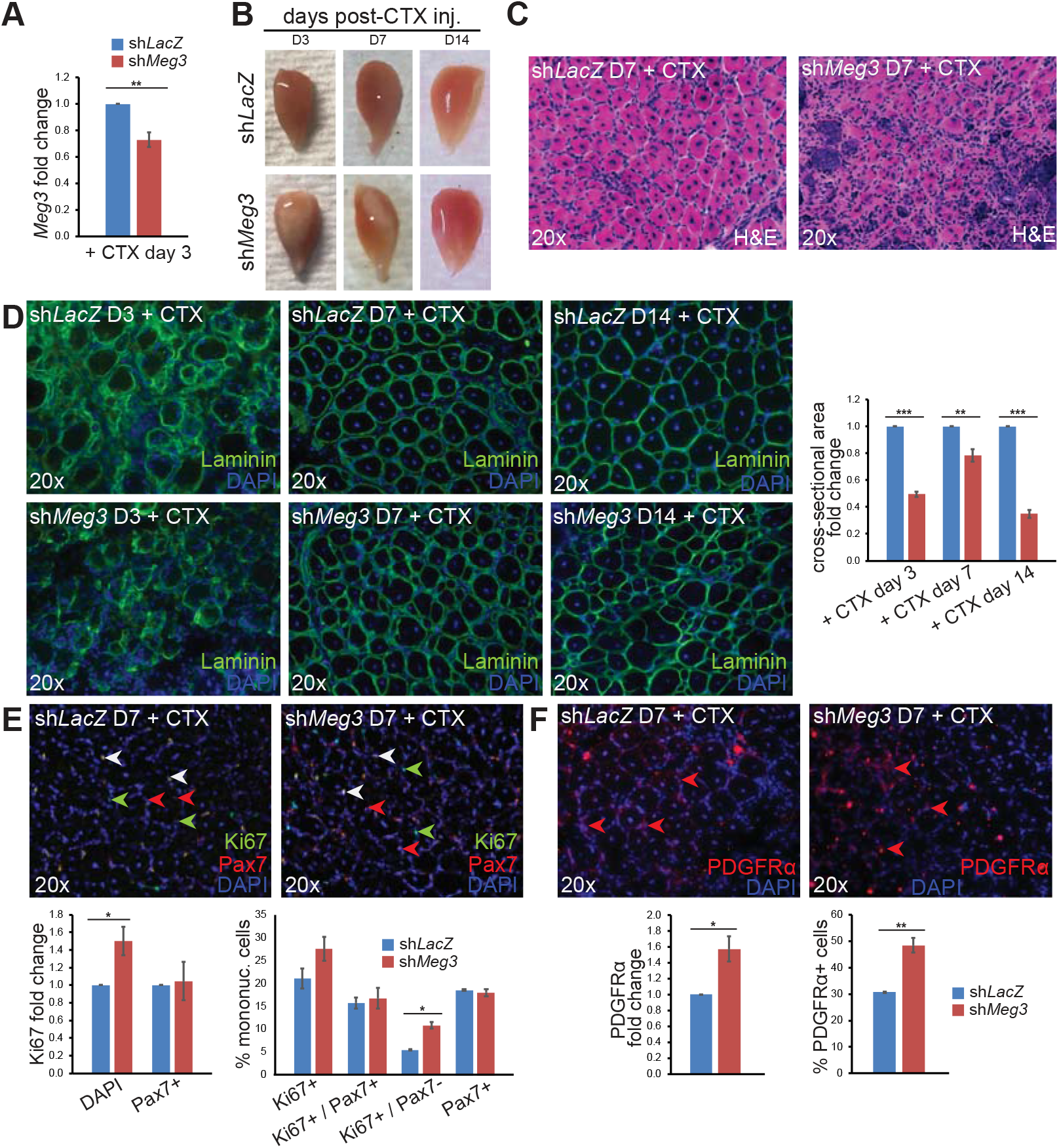
*Meg3* knockdown impairs injury-induced skeletal muscle regeneration. **A)** qPCR expression profiling indicated reduced *Meg3* expression in TA muscles co-injected with sh*Meg3* adenovirus (n=3). **B)** Whole mount morphology of regenerating muscles co-injected with adeno-sh*LacZ* (top) or adeno-sh*Meg3*. **C)** Hematoxylin and eosin (H&E) staining of muscle sections. **D)** Cross-sectional area (CSA) of laminin-ensheathed regenerating myofibers was measured for days 3, 7, and 14 post-CTX injury. sh*Meg3* muscle displayed reduced CSA for all time points surveyed. **E)** Immunofluorescence quantification indicated sh*Meg3* TA sections harbor increased Ki67 signal (left bar graph). Co-staining for Pax7 indicated no change in satellite-cell specific Ki67 signal (left graph) and no change in satellite cell abundance (white, red arrowheads). Marker quantification revealed an increase in proliferating cells lacking Pax7 co-stain (green arrowheads). **F**) Immunofluorescence quantification indicated an increase in PDGFR signal, as well as increased abundance of PDGFRα+ cells (red arrowheads).

Whole-mounts of regenerating TA muscle treated with sh*Meg3* adenovirus exhibited more pronounced necrotic regions at day 3 and 7 post-injury that diminished by day 14 (Fig. 8B). Regenerating sh*Meg3* muscle sections also harbored regions with intense basophilic stain at a higher frequency than observed in sh*LacZ* control TA muscle (Fig. 8C). To evaluate for changes in cross-sectional area (CSA), an indicator of myoblast fusion, injured TA muscle sections were analyzed by laminin immunofluorescence. While *Meg3* levels increased in injured sh*Meg3* TA muscle by 7 days post-injury (data not shown), the CSA of sh*Meg3* myofibers was not only significantly reduced in sh*Meg3*-treated TA muscle at day 3, but also at day 7 and 14 post-injury (Fig. 8D). These findings suggest that inhibition of *Meg3* at the onset of injury, when muscle and non-muscle precursors are activated, had long-lasting detrimental effects on muscle regeneration.

### Increased proliferation of mesenchymal stromal but not satellite cells in injured sh*Meg3* muscle

Given that *Meg3* has well-documented roles modulating proliferation in a variety of cell types, and the proliferation defect observed in sh*Meg*3 C2C12 myoblasts, we considered the possibility that *Meg3* modulates this process in regenerating muscle. To this end, proliferation in muscle stem cells was examined by immunofluorescence using Ki67, a marker of actively cycling cells, and Pax7, a muscle stem cell (satellite cell) marker. Quantification of Ki67 revealed a significant increase in protein signal, but this elevated expression was independent of satellite cells (Fig. 8E, lower left bar graph). Consistent with this observation, there was a significant increase in the number of Ki67+/Pax7-, but not Ki67+/Pax7+ mononucleated cells (Fig. 8E, lower left graph). Moreover, the total number of Pax7+ cells did not change in regenerating sh*Meg*3-treated muscle at day 7 post-injury (Fig. 8E, lower left). These data reveal that elevated cell cycle activity in regenerating *Meg3*-deficient muscle is attributable to a non-satellite cell population.

To define this non-satellite cell population, we subsequently examined expression of PDGFRα, a marker for limb muscle mesenchymal stromal cells – also known as fibroadipogenic progenitors (FAPs) (Joe *et al.* 2010) – that express *Meg3* lncRNA (Fig. 1A). As shown in Fig. 8F, a marked increase in expression and number of PDGFRα+ cells was observed in sh*Meg*3 relative to sh*lacZ* control muscle. These data indicate that *Meg3* knockdown stimulates mesenchymal stromal cell expansion rather than changes in the abundance of satellite cells, and suggest that reduced myofiber CSA is not a consequence of reduced satellite cell number, but rather defects in satellite cells unrelated to proliferation.

### Enhanced mesenchymal-like properties in *Meg3*-depleted regenerating muscle

To precisely define the molecular mechanisms of impaired myofiber formation in *Meg3*-deficient regenerating muscle, RNA-seq was performed on TA muscle tissue 7 days post-injury. The dysregulated transcriptome of injured sh*Meg3*-treated muscle revealed significant enrichment of genes involved in EMT (Hallmark) and TGFβ signaling (Hallmark and IPA), similar to that observed in *Meg3*-deficient C2C12 myoblasts (Fig. 9A - D).

**Figure 9.**
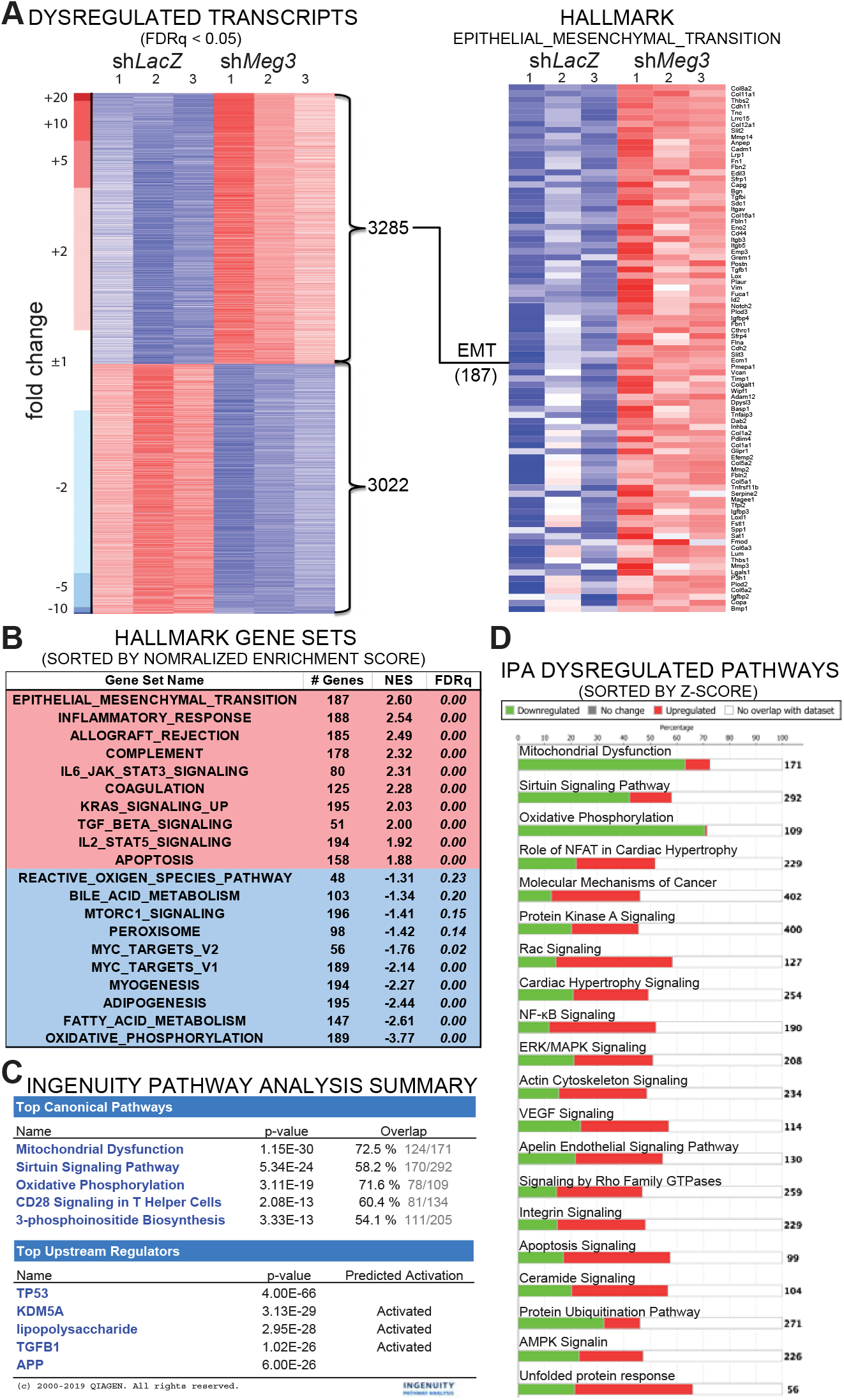
EMT and TGFβ signaling are among the top dysregulated pathways in regenerating *Meg3* knockdown muscle. **A)** Heatmap of 6,307 dysregulated transcripts, with 3,285 upregulated (red) and 3,022 downregulated (blue). Broad GSEA Hallmark analysis indicated coherent upregulation of 187 well-characterized EMT gene markers. **B)** Table conveying the top 10 up (red) and down (blue)-regulated Hallmark biological states and cellular processes, as indicated by normalized enrichment score (NES). EMT was the top upregulated gene sets. **C)** Summary output from Quiagen Ingenuity Pathway Analysis (IPA) software performed with significantly dysregulated transcripts (p<0.05). IPA lists TGFβ1 among Top Upstream Regulators, and lists various metabolic pathways among the Top Canonical Pathways. **D)** Stacked bar graphs indicate proportions of upregulated (red) and downregulated (green) genes that comprise the top 20 dysregulated biological pathways, which include metabolic pathways (Mitochondrial dysfunction, Sirtuin signaling, Oxidative phosphorylation, Protein Kinase A Signaling), RhoA-related pathways (Rac signaling, Signaling by Rho Family GTPases) and EMT-related pathways (Molecular Mechanisms of Cancer, ERK/MAPK signaling, Actin Cytoskeleton Signaling, Integrin signaling).

Based on these results, we examined for cadherin switching, and confirmed that *N-cadherin* transcripts were upregulated in sh*Meg3* muscle 7 days post-injury (Fig. 10A left graph). In contrast, *E-cadherin* transcripts were unchanged, and protein signal was undetectable (data not shown). Detailed examination of N-cadherin protein at the cellular level revealed that its expression was largely restricted to Pax7+ cells. Though N-cadherin intensity per satellite cell was unchanged (Fig. 10A right graph), the percentage of satellite cells expressing N-cadherin was significantly increased in *Meg3*-deficient muscle (Fig. 10A middle graph). These data indicate an enrichment of mesenchymal adhesion properties in satellite cells in regenerating sh*Meg3*-treated muscle even in the absence of a *bona fide* switch.

**Figure 10.**
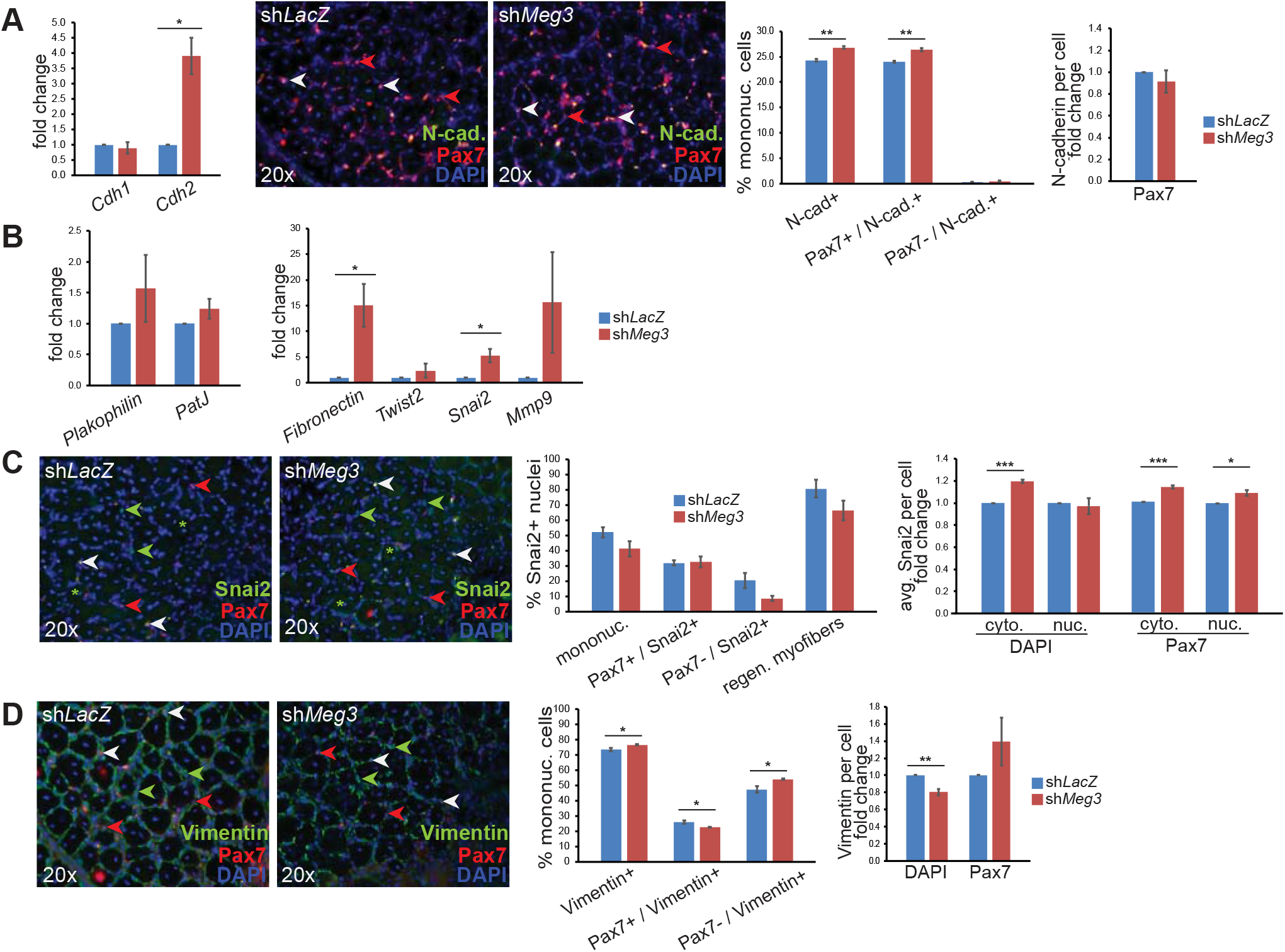
Enhanced mesenchymal character in *Meg3* knockdown regenerating muscle microenvironment. **A)** Expression profiling by qPCR indicated no change in *E-cadherin*, while day 7 sh*Meg3* muscle was enriched for *N-cadherin* transcripts (n=3). Immunofluorescence revealed that, while numerous satellite cells were not N-cad+ (red arrowheads), N-cad signal was largely restricted to Pax7+ satellite cells in regenerating muscle (white arrowheads). Cell quantification (% mononuc. cells) revealed that sh*Meg3* muscle harbored increased abundance of N-cadherin+ satellite cells, whereas levels of N-cadherin per cell was unchanged (n=3). **B)** qPCR expression profiling indicated no change in epithelial markers *Plakophilin* and *PatJ*, but mesenchymal markers *Fibronectin* and *Snai2* were significantly upregulated in sh*Meg3* muscle (n=3). **C)** Immunofluorescence revealed the presence of satellite cells lacking Snai2 (red arrowheads), Snai2+ satellite cells (white arrowheads), Snai2+ non-satellite cells (green arrowheads), and Snai2+ nuclei in regenerating myofibers (green asterisks). Quantification of Snai2+ cells indicated no change in the occurance of Snai2+ nuclei (bar graph, % Snai2+ nuclei). Generalized analysis (DAPI) indicated significant upregulation of cytoplasmic Snai2 signal per cell, but no change nuclear intensity; Snai2 signal in satellite cells (Pax7+) was increased for both cytoplasmic and nuclear compartments (n=3). **D)** Immunofluorescence revealed the presence of satellite cells lacking Vimentin (red arrowheads), Vimentin+ satellite cells (white arrowheads), and Vimentin+ non-satellite cells (green arrowheads). Cell quantification (% mononuc. cells) revealed increased abundance of Vimentin+ mononucleated cells that were not Pax7+, and that Pax7+/Vimentin+ cells were reduced in sh*Meg3* muscle. Vimentin signal per cell was downregulated in mononucleated cells (bottom right bar graph), which may reflect sh*Meg3*-specifici differences in Vimentin+ cell morphology (green channel) (n=3).

To further analyze EMT in regenerating muscle, expression of additional epithelial and mesenchymal markers was examined. Epithelial marker genes *Plakophilin* and *PatJ* were unaffected by sh*Meg3* treatment, but mesenchymal markers *Fibronectin* and *Snai2* were significantly increased (Fig. 10B). To delve deeper into the increased mesenchymal marker expression observed in sh*Meg3* muscle, we analyzed protein localization of the EMT transcription factor Snai2, which is regulated, in part, through nucleo-cytoplasmic shuttling (Lamoiulle et al 2014). Snai2 was detected in satellite cells, Pax7-mononucleated cells, and regenerating myofibers, and the proportions of these Snai2+ cells did not change with sh*Meg3* treatment (Fig. 10C, left graph). However, while all Snai2+ cells had increased cytoplasmic Snai2 (Fig. 10C, right graph, DAPI), only satellite cells displayed significantly more intense nuclear Snai2 signal compared to sh*LacZ* controls (Fig. 10C, right graph, Pax7). Although multiple cell types displayed aberrant accumulation of Snai2 in the cytoplasm, its increased nuclear localization in satellite cells is consistent with upregulation of other mesenchymal genes, and strongly suggests an increased ability to activate EMT in these progenitors.

Vimentin expression was analyzed and was detected in numerous mononucleated cells, including a subset of satellite cells, in the regenerating muscle microenvironment. While the overall percentage of vimentin+ mononucleated cells was increased, this net increase can be attributed to Pax7-cells, as vimentin+ satellite cells were reduced (Fig. 10D, left graph). Interestingly, with the exception of satellite cells, there was less vimentin expressed per cell in sh*Meg3* muscle reflecting an overall reduction in its synthesis (Fig. 10D, right graph). Taken together, *Meg3* inhibition predominantly impacted vimentin expression in cells that are not of satellite cell origin.

Overall, these data reveal that knockdown of *Meg*3 in the context of cardiotoxin-induced muscle injury dramatically remodeled the regenerating tissue microenvironment by triggering aberrant EMT activation in satellite cells and - to a lesser extent - non-satellite cells.

## Discussion

Our findings reveal a key role for the lncRNA *Meg3* in myoblast plasticity and highlight the importance of proper regulation of EMT and TGFβ signaling for myogenic differentiation. Chronic depletion of *Meg3* in C2C12 myoblasts resulted in an enhanced mesenchymal cell state, marked by imbalance of epithelial and mesenchymal genes, which diminished growth and differentiation. Moreover, *Meg3* inhibition in skeletal muscle injury severely impaired myofiber formation, increased mesenchymal gene expression in satellite cells, and resulted in abnormal mesenchymal stromal cell expansion, radically altering the cellular composition in the regenerating microenvironment. In both models, transcriptomic profiling indicated that *Meg3* regulates EMT through repression of TGFβ signaling, and inhibition of TGFβR1 - or its downstream effectors RhoA and p38MAPK - was sufficient to restore myogenic differentiation *in vitro*.

Like many imprinted genes, expression of the *Dlk1-Dio3* locus correlates with changes in cell state (Stadtfeld *et al.* 2010; Liu *et al.* 2010; Haga and Phinney 2012; Qian *et al.* 2016), and *Meg3* has emerged as a major driver of these adaptations. *Meg3* has been shown to mediate stem cell pluripotency and regulate the differentiation potential of progenitor cells belonging to diverse lineages (Kaneko *et al.* 2014; Li *et al.* 2017; Gokey *et al.* 2018; Yen Y-P *et al.* 2018). Moreover, its inhibition activates EMT to promote mesenchymal behavior in cancer cells (Terashima *et al.* 2017; Deng *et al.* 2018; Yu *et al.* 2018). Our findings intersect with these reported functions and demonstrate that *Meg3* is required to prevent inappropriate EMT activation in myoblasts.

While EMT is a requisite step for embryonic muscle development, these acquired mesenchymal traits must be kept in check to ensure successful cell state transitions for differentiation. Loss of this checkpoint control in *Meg3*-depleted myoblasts resulted in increased TGFβ and Rho activity, which are not only drivers of EMT, but also potent inhibitors of myogenesis. Enrichment of N-cadherin, another major mediator of EMT, likely exacerbated the differentiation defects in *Meg3-*depleted myoblasts, as N-cadherin has been shown to trigger RhoA activation (Lovett *et al.* 2006), and excessive activity of RhoA compromises cell-cell adhesion and fusion competence in myogenic cells (Lu *et al.* 2010). Furthermore, Snai2 (Slug) was enriched in *Meg3*-depleted myoblasts and activated satellite cells, and this key transcriptional regulator not only reinforces EMT via suppression of epithelial gene programs, but has also been shown to obstruct myogenic differentiation via occupancy of MyoD E-box enhancers (Hajra *et al.* 2002; Soleimani *et al.* 2012). Consequently, our experiments revealed that ineffective EMT repression in *Meg3*-depleted myoblasts and injured skeletal muscle ultimately impaired myotube formation, which underscores an important modulatory role for *Meg3* in myoblast plasticity.

Given that aberrant TGFβ activation contributed to the EMT and differentiation defects in *Meg3-*depleted myoblasts, *Meg3*-mediated epigenetic repression of TGFβ-related loci could be a nodal regulatory mechanism for maintaining proper cell state. Similar to how *Meg3* interacts with Ezh2 to repress TGFβ signaling in breast cancer cells (Mondal *et al.* 2015), it is conceivable that *Meg3* coordinates PRC2 function with TGFβ activity to fortify myogenic cell character. PRC2 is an important epigenetic regulator of muscle stem cell identity and myogenic differentiation (Caretti *et al.* 2004; Juan *et al.* 2011; Woodhouse *et al.* 2013), yet there is limited information on how its recruitment is regulated in muscle. In support of this notion is the observation that *Meg3* expression overlaps with that of Ezh2 in satellite cells (Wust *et al.* 2018; Juan *et al.* 2011), and our results demonstrate an interaction between *Meg3* and Ezh2 in proliferating C2C12 myoblasts. Thus, *Meg3* represents a strong candidate for PRC2 recruitment specificity in myoblasts directing this epigenetic factor to generate a chromatin landscape that both maintains identity and primes these cells for differentiation.

While *Meg3* is classified as a tumor suppressor, the reduced proliferation caused by its inhibition in myoblasts *in vitro* seems to reflect the context-dependent functions of *Meg3*. Its inhibition resulted in an upregulation in N-cadherin, which has been shown to suppress cell cycle re-entry in quiescent satellite cells (Goel *et al.* 2017). Furthermore, Ezh2 facilitates satellite cell expansion (Juan *et al.* 2011; Woodhouse *et al.* 2013), thus perturbations in its function in *Meg3*-depleted C2C12 myoblasts could also impair proliferation in these cells. Surprisingly, myoblast proliferation was not affected in regenerating sh*Meg3*-treated muscle even though aberrant stimulation of TGFβ is an impediment to satellite cell activation and proliferation (Rathbone *et al.* 2011). Instead, mesenchymal stromal cell (MSC) expansion was observed. Considering MSCs are the only Pax7-interstitial cell population expressing appreciable levels of *Meg3*, we cannot exclude the possibility that *Meg3* knockdown in MSCs triggered this expansion, suggesting a differential role for *Meg3* in interstitial cell growth. It is noteworthy that mesenchymal stromal cells facilitate regeneration (Wosczyna *et al.* 2019), but abnormal accumulation of these cells is associated with fibrosis in dystrophic muscle (Ito *et al.* 2013; Malecova *et al.* 2018), which could hinder proper allocation of satellite cells during the regenerative process. Future single cell analysis of mononucleated cell populations may provide a deeper understanding of the signaling mechanisms through which *Meg3* regulates muscle and non-muscle cell types in regenerating muscle.

Activation of TGFβ is a pathological hallmark for both aged and diseased muscle (Burks and Cohn 2011). Indeed, the phenotype of regenerating sh*Meg3*-treated muscle conspicuously resembles age-related muscle defects (Beggs *et al.* 2004; Lukjanenko *et al.* 2019). Recently, a subset of *Dlk1-Dio3* miRNAs have been shown to suppress age-related atrophy (Shin *et al.* 2020), and these ncRNAs – including *Meg3* – are markedly downregulated in aged muscle (Mikovic *et al.* 2018). Given that aged satellite cells accumulate epigenetic abnormalities that suppress their ability to efficiently regenerate muscle (Liu et al 2013; Sousa-Victor et al, 2014), we postulate that age-related *Meg3* downregulation in muscle induces a reprogramming of the epigenetic landscape that is more sensitized to pathological TGFβ and EMT signaling. Thus, *Meg3* represents a promising avenue of investigation for therapeutics targeting the deleterious physiology of muscle wasting and dysfunction.

## Materials and methods

### Cell culture

C2C12 myoblasts were grown and induced to differentiate as described previously (Huang et al 2006). For stable knockdown, C2C12 myoblasts were seeded at 20-30% confluence in 10cm format (CELLSTAR) and forward-transfection with Fugene-6 (Promega) according to manufacturer’s protocol with the following plasmids: 9μg shRNA plasmid (pENTRU6-sh*LacZ* or pENTRU6-sh*Meg3*) and 3μg pCDNA3 empty vector. For G418 selection, cells were grown in antibiotic-free growth medium supplemented with Geneticin (0.18mg/mL, Gibco), which was changed daily for 12 days. For heterogeneous studies, cells were used for experiments 48 hours following selection. For rescue/overexpression experiments, plates were transfected with 9μg overexpression plasmid (pShuttleCMV-β-gal or pShuttleCMV-MEG3), 3μg shRNA plasmid (pENTRU6-sh*LacZ* or pENTR U6-sh*Meg3*), and 1μg resistance plasmid (pCDNA3 empty vector).

### RNA immunoprecipitation

60 plates of C2C12 myoblast cell pellets were collected on ice, and dounce homogenized in RNase-free AT buffer (20% glycerol, 1mM EDTA, 0.15M NaCl, 20nM HEPES pH 7.7, DTT, Protease inhibitors, RNAsin). Equal volumes of cleared lysate were added to 45μL Protein G beads (Santa Cruz SC2002) preincubated with 0.5μL Ezh2 antibody (Millipore 07-689) or normal mouse IgG (Santa Cruz SC2025). Following overnight nutation at 4°C, 45μL supernatant was collected for normalization control, and immunoprecipitates were washed four times with AT buffer before resuspending bead conjugates in Trizol for RNA purification.

### cDNA synthesis and qPCR

RNA from C2C12 myoblasts or TA muscle was extracted with Trizol Reagent (Ambion) and used according to manufacturer’s protocol. RNA template was converted to cDNA using M-MLV reverse transcriptase, random oligo dT primers, and RNAsin according to manufacturer’s protocol (Promega). Quantitative RT-PCR was performed in triplicate wells using Power SYBR® Green Master Mix (Applied Biosystems) with the 7900HT Sequence Detection System (Applied Biosystems). Primers used for all qRT-PCR analyses are listed in Supplemental Table S1.

### Plasmids

For overexpression, human *MEG3* cDNA was PCR amplified from the pCI-Meg3 (Addgene Plasmid #44727) using NEB Q5 high-fidelity polymerase, and cloned into pShuttleCMV vector (Agilent AdEasy Adenoviral Vector System). For knockdown, the sh*Meg3* target sequence was derived from 2015 Mondal *et al.*, and double-stranded RNA insert was ligated to U6-pENTR shuttle vector (BLOCK-iT^TM^ U6 RNAi Entry Vector Kit, Invitrogen). Construction of sh*LacZ* was described previously (Ewen et al., 2011). Plasmids were grown in DH5α cells and purified via midi-prep columns (Machery-Nagel Nucleobond).

### Adenovirus

To generate Adenoviruses used for muscle regeneration, pENTR U6-shRNA plasmids were ligated to pAd plasmid and transfected into HEK293A cells. Following plaque formation, crude viral lysates were precipitated and purified by CsCl purification and dialysis. Virus was tittered by serial dilution and plaque formation.

### Muscle injury

*Tibialis anterior* (TA) muscle of Swiss-Webster mice (female, 25 grams, Taconic) were injected with 10 μM cardiotoxin (*Naja nigricollis,* EMD chemicals) in saline solution containing 5×10^10^ pfus adenovirus, and administered as 50 μL injections per TA muscle. An additional round of virus was injected 24 hours post-CTX injury, and muscles were harvested 3, 7, and 14 days post-injury.

### Inhibitors and growth factors

All chemical inhibitors were resuspended using the vendor’s specifications, and added to cells at the indicated concentration by diluting into culture medium for day of seeding (day −1), and differentiation (day 0), with 1, 2, 3). Media was replaced daily (with one wash) to ensure additives were fresh. Final concentrations: LY2157299 (10μM, Sigma), Y-27632 (40μM Sigma), SB203580 (10μM, Calbiochem), BMP4 (50ng/μL Life Technologies).

### Immunofluorescence

C2C12 myoblasts and transverse TA muscle cryosections (16μm) were post-fixed with 4% para-formaldehyde and blocked with PBS solution containing 0.3% Triton-X 100 and 3% normal Donkey serum (Sigma). Antibody dilution buffer consisted of PBS containing 1% BSA and 0.3% Triton-X 100, and samples were immunostained as described in Kanisiack et al. (2009). Primary antibodies: MYH4 (1:200, Proteintech 20140-1-AP), α-actinin (1:1000, Sigma A7811), BrdU (1:500, Invitrogen MoBU-1), Ki67 (1:2000, Abcam ab15580), Laminin (1:2000, Sigma L9393), Pax7 (1:200, DSHB PAX7), PDGFRα (1:200, Proteintech 60045-1-Ig), Cdh1 (1:500, Abcam ab40772), Cdh2 (1:500, Abcam ab76011), Vimentin (1:1000, Abcam ab92547). Secondary antibodies: donkey anti-mouse Alexa-Fluor 488 IgG (1:2000, Invitrogen), donkey anti-mouse Alexa-Fluor 568 IgG (1:2000, Invitrogen), donkey anti-rabbit Alexa-Fluor 488 IgG (1:2000, Invitrogen), donkey anti-rabbit Alexa-Fluor 555 IgG (1:2000, Invitrogen). All samples were mounted with DAPI Vectashield mounting medium. For Pax7 immunostaining, immunostaining was modified to follow protocol for Mouse on Mouse Immunodetection kit (Abcam). For BrdU, proliferating C2C12 myoblasts were pulsed with 10μM BrdU (BD Pharmingen) for 30 minutes followed by fixation, incubated with 2.5M HCl solution at 37°C for 20 minutes, and washed four times with PBS prior to proceeding to immunostaining.

### MitoTrackerCMXRos

MitoTracker reagent was resuspended by manufacturer’s protocol in DMSO, and added to C2C12 cultures to a final concentration of 50μM. Following 30 minute incubation, cells were fixed and subjected to immunostaining.

### Image acquisition

BrdU and laminin stained images were acquired using Olympus DSU All other immunofluorescent and brightfield images were captured using Nikon Eclipse NiE (BU Proteomics and Imaging Core).

### Image analyses

Fusion index was quantified using ImageJ Cell Counter Plugin. Immunofluorescence was measured on ImageJ with Intensity Ratio Nuclei Cytoplasm Tool (http://dev.mri.cnrs.fr/projects/imagej-macros/wiki/Intensity_Ratio_Nuclei_Cytoplasm_Tool). Scratch wound assay scratch areas were measured with the MRI Wound Healing Tool (https://github.com/MontpellierRessourcesImagerie/imagej_macros_and_scripts/wiki/Wound-Healing-Tool). Cross-sectional area (CSA) of skeletal muscle was assessed in laminin-stained sections using ImageJ threshold, binary, watershed functions and area tracing tools.

### RNA sequencing

RNA samples corresponding to day 3 C2C12 differentiation (n=3 sh*LacZ*, n=3 sh*Meg3*) and day 7 injured TA muscle (n=3 sh*LacZ*, n=3 sh*Meg3*) were submitted to the Boston University Microarray and Sequencing Resource Core for library prep and Illumina sequencing.

### Western blot analysis

C2C12 cells were lysed by dounce-homogenization in the presence of ELB with DTT and protease inhibitors. Protein concentration was determined by Bradford (Bio-Rad). Western blots were performed as previously described (McCalmon et al., 2010), with the following modifications: primary antibodies were incubated overnight in BLOTTO solution at 4°C; subsequent washes were gradually transitioned to PBS for 1 hour Licor dye incubation; membranes were shielded from light, and pressed dry with Whatman paper prior to image acquisition (Azure Sapphire Biomolecular Imager). β-tubulin (1:5000, Cell Signaling technology mAb #3873), MF20 (0.2 μg/mL, DSHB MF-20 supernatant), Cdh1 (1:1000, Abcam ab40772), Cdh2 (1:1000, Abcam ab76011), Vimentin (1:2000, Abcam ab92547). Secondary dyes: Rabbit 800 (1:2000, Licor 926-32211), Mouse 680 (1:2000, Licor 926-68070)

### Statistical Analysis

All numerical quantification is representative of the mean ± S.E.M. of at least three independently performed experiments. Statistically significant differences between two populations of data were determined using the student’s T-test. P-values of ≤0.05 were considered to be statistically significant. * = p<0.05, ** = p<0.01, *** = p<0.001, for student’s t-test. Bar graph data are presented as mean ± std. error.

## Acknowledgments

We are grateful to the BU Sequencing Core for RNA sequencing and Adam Gower for bioinformatics analysis. We also thank Andrew Emili (Director, Center for Systems Biology, BU School of Medicine), Angela Ho (BU Dept of Biology) for providing us with additional reagents and advice, and Tarik Zahr and Roxanna Altus for excellent technical assistance.

## Data availability

Single Cell RNAseq data were obtained from the FACS data visualization, provided by the Chan-Zuckerburg Initiative BioHub (https://tabula-muris.ds.czbiohub.org/). Permission to use BioHub data in this manuscript’s Figure 1 was granted by Tabula Muris corresponding author Spyros Darmanis. http://microarray.bu.edu/~msrdata/2018-06-21_Naya/2018-06-21_Naya_analysis.xlsx http://microarray.bu.edu/~msrdata/2018-06-21_Naya/2018-06-21_Naya_GSEA.xlsx

---

**Supplemental Figure 1.**
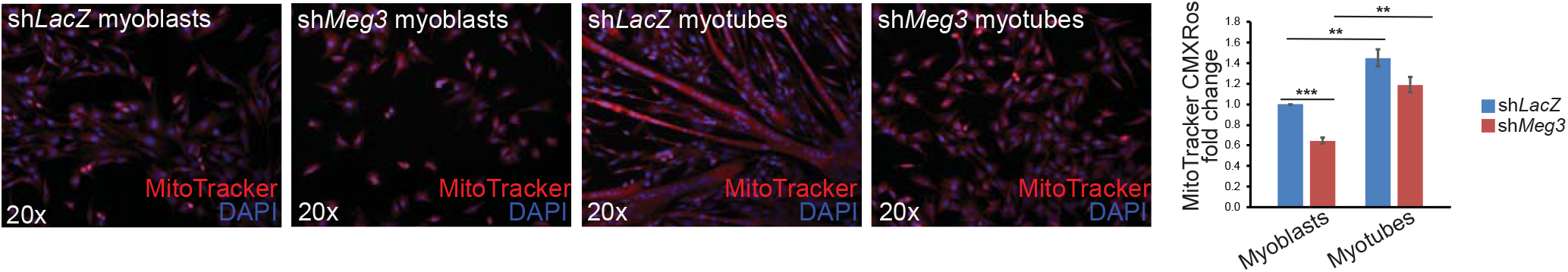
sh*Meg3* myoblasts display reduced mitochondrial mass. C2C12 myoblasts and myotubes were pulsed with MitoTracker CMXRos for 40 minutes, and co-stained with α-actinin. Quantification of Mitotracker (restricted to α-actinin+ cells) indicated reduced mitochondrial signal in sh*Meg3* myoblasts, but not myotubes (n=3). Both treatment groups displayed increased MitoTracker signal with differentiation.

**Supplemental Figure 2.**
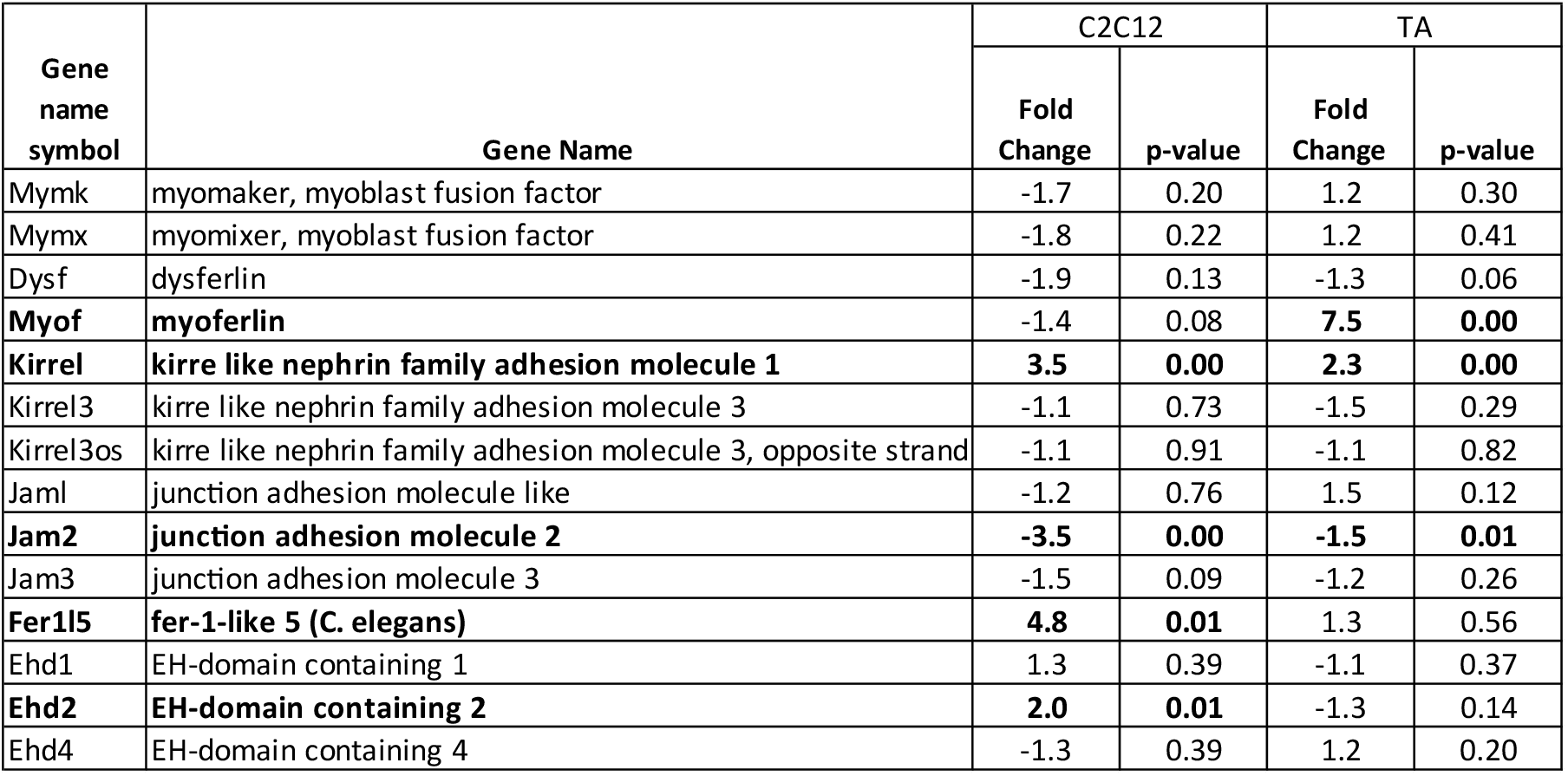
Myogenic fusion transcripts are not downregulated in sh*Meg3* myotubes. RNAseq data indicated that transcripts of master regulators of myogenic fusion, notably *Myomaker* and *Myomixer*, were not significantly downregulated in sh*Meg3* myotubes.

**Supplemental Figure 3.**
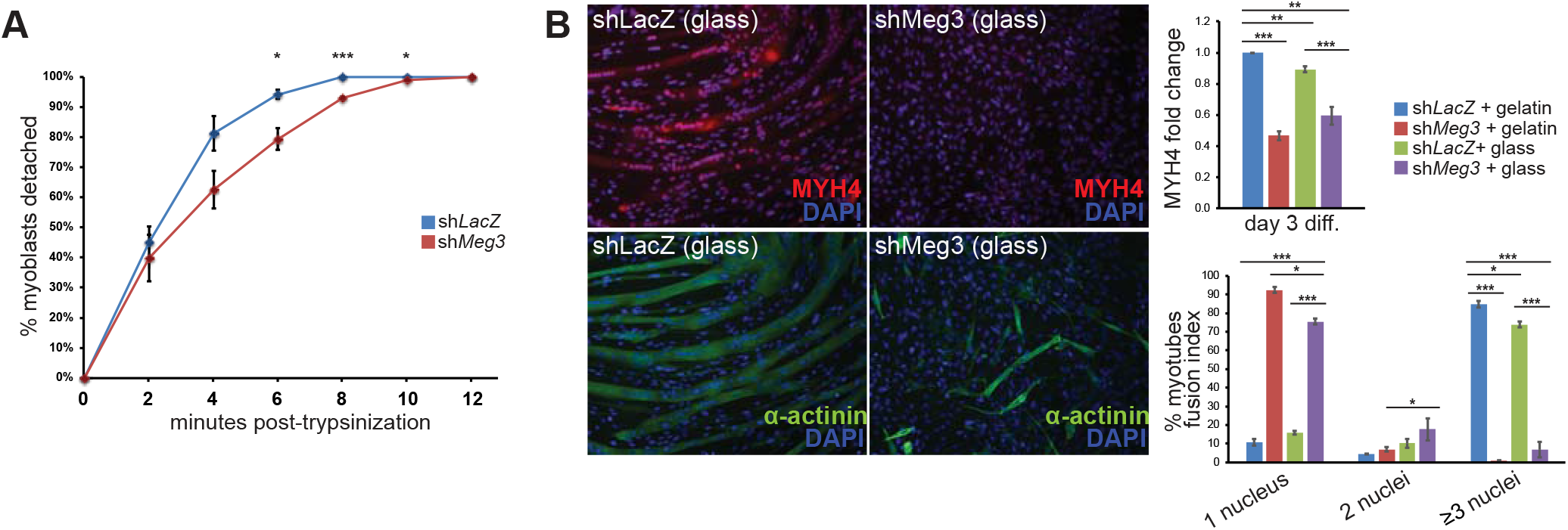
sh*Meg3* myoblasts require longer trypsinzation, and depriving cells of surface substrate had modest effects on fusion. **A)** Quantification of cells recovered over time during trypsinization revelaed that sh*Meg3* myoblasts require significantly more time to trypsinize to completion (n=3). **B)** When differentiated upon a substrate-deprived surface (glass), sh*Meg3* myoblasts had no change on MYH4 expression (n=3) or fusion of cells with ≥3 nuclei, but did exhibit reduced quantities of myotubes with 1- and 2- nuclei relative to sh*Meg3* controls differentiated on 0.1% gelatin (n=3). Differentiation atop glass also significantly reduced sh*LacZ* MYH4 signal (n=3) and quantity of myotubes with ≥3 nuclei.

**Supplemental Figure 4.**
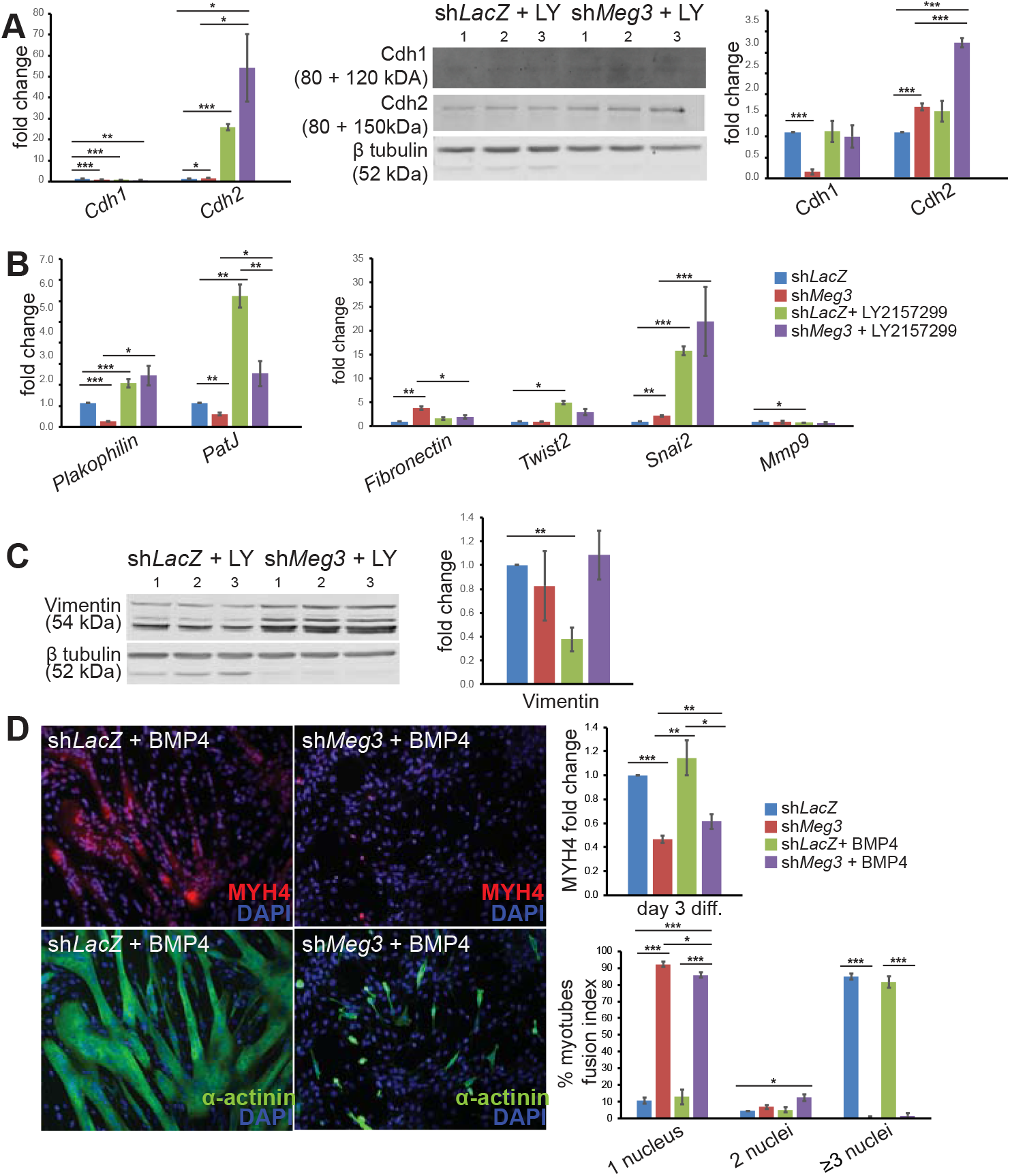
TGFβR1 inhibition results in dynamic EMT marker expression, and BMP4 stimulation is not sufficient for sh*Meg3* rescue. **A)** qPCR indicated that LY2157299 (LY) treatment resulted in reduced *E-cadherin* (*Cdh1*) transcripts regardless of sh*RNA* treatment, with simultaneous upregulation of *N-cadherin* (*Cdh2*) transcripts (n=3). Western blot revealed modest Cdh1 band detection, and quantification of β-tubulin-normalized signal revealed that LY-treatment restored Cdh1 levels to sh*Meg3* myotubes, while Cdh1 in sh*LacZ* myotubes remained unchanged. While LY treatment did not change Cdh2 expression in sh*LacZ* myotubes, LY-treatment enhanced Cdh2 signal in sh*Meg3* myotubes. **B)** qPCR profiling indicated upregulation of epithelial transcripts *Plakophilin* and *PatJ* regardless of sh*RNA* background (n=3). *Fibronectin* transcript levels returned to normal levels in LY-treated sh*Meg3* myotubes. LY treatment intensified upregulation of *Snai2* transcripts in sh*Meg3* cells, but did not affect *Twist2* or *Mmp9* levels relative to sh*Meg3* myotubes. sh*LacZ* + LY myotubes displayed reduced *Mmp9*, with simultaneous upregulation of *Twist2* when compared to untreated sh*LacZ* cells (n=3). **C)** Western blot quantification of Vimentin suggests that LY treatment reduced Vimentin expression in sh*LacZ* controls, but did not change Vimentin expression in sh*Meg3* myotubes (n=3). **D)** Myoblasts pre-treated with 5ng/mL BMP4 (BMP) were subjected to differentiation, and examined for changes in MYH4 expression and fusion index. BMP4 treated sh*Meg3* myotubes had improved MYH4 expression (n=3), reduced mononucleated myotubes, and improved 2-cell fusion, but not ≥3 nuclei fusion.

## References

Beggs ML, Nagarajan R, Taylor-Jones JM, Nolen G, Macnicol M, Peterson CA. (2004) Alterations in the TGFbeta signaling pathway in myogenic progenitors with age. Aging Cell. 3(6):353–61.

Benetatos L, Hatzimichael E, Londin E, Vartholomatos G, Loher P, Rigoutsos I, Briasoulis E. (2013) The microRNAs within the DLK1-DIO3 genomic region: involvement in disease pathogenesis. Cell Mol Life Sci. 70(5):795–814.

Benetatos L, Vartholomatos G, Hatzimichael E (2014). DLK1-DIO3 imprinted cluster in induced pluripotency: landscape in the mist. Cell Mol Life Sci. 71(22):4421–30.

Buckingham M, Rigby PW(2014). Gene regulatory networks and transcriptional mechanisms that control myogenesis. Dev Cell. 28(3):225–38.

Burks TN, Cohn RD.(2011) Role of TGF-β signaling in inherited and acquired myopathies. Skelet Muscle. 1(1):19.

Campbell K, Casanova J.(2016) A common framework for EMT and collective cell migration. Development. 143(23):4291–4300.

Caretti G, Di Padova M, Micales B, Lyons GE, Sartorelli V.(2004) The Polycomb Ezh2 methyltransferase regulates muscle gene expression and skeletal muscle differentiation. Genes Dev. 18(21):2627–38.

Castel D, Baghdadi MB, Mella S, Gayraud-Morel B, Marty V, Cavaillé J, Antoniewski C, Tajbakhsh S (2018). Small-RNA sequencing identifies dynamic microRNA deregulation during skeletal muscle lineage progression. Sci Rep. 8(1):4208.

Chal J, Pourquié O (2017). Making muscle: skeletal myogenesis in vivo and in vitro. Development. 144(12):2104–2122.

da Rocha ST, Edwards CA, Ito M, Ogata T, Ferguson-Smith AC (2008). Genomic imprinting at the mammalian Dlk1-Dio3 domain. Trends Genet. 24(6):306–16.

Das PP, Hendrix DA, Apostolou E, Buchner AH, Canver MC, Orkin S (2015). PRC2 Is Required to Maintain Expression of the Maternal Gtl2-Rian-Mirg Locus by Preventing De Novo DNA Methylation in Mouse Embryonic Stem Cells. Cell Rep. 12(9):1456–70.

Deng R, Fan FY, Yi H, Liu F, He GC, Sun HP, Su Y. (2018). MEG3 affects the progression and chemoresistance of T-cell lymphoblastic lymphoma by suppressing epithelial-mesenchymal transition via the PI3K/mTOR pathway. J Cell Biochem.

Dill, T.L. and Naya, F.J. (2018) A Hearty Dose of Noncoding RNAs: The Imprinted DLK1-DIO3 Locus in Cardiac Development and Disease. J Cardiovasc Dev Dis. 5(3).

Eisenberg, I., Eran, A., Nishino, I., Moggio, M., Lamperti, C., Amato, A.A., Lidov, H.G., Kang, P.B., North, K.N., Mitrani-Rosenbaum, S., et al. (2007). Distinctive patterns of microRNA expression in primary muscular disorders. Proc. Nat. Acad. Sci. USA 104(43), 17016–21.

Ewen EP, Snyder CM, Wilson M, Desjardins D, Naya FJ. (2011) The Mef2A transcription factor coordinately regulates a costamere gene program in cardiac muscle. J Biol Chem. 286(34):29644–53.

Gao YQ, Chen X, Wang P, Lu L, Zhao W, Chen C, Chen CP, Tao T, Sun J, Zheng YY, Du J, Li CJ, Gan ZJ, Gao X, Chen HQ, Zhu MS. (2015) Regulation of DLK1 by the maternally expressed miR-379/miR-544 cluster may underlie callipyge polar overdominance inheritance. Proc Natl Acad Sci U S A. 112(44):13627–32.

Goel AJ, Rieder MK, Arnold HH, Radice GL, Krauss RS. (2017) Niche Cadherins Control the Quiescence-to-Activation Transition in Muscle Stem Cells. Cell Rep. 21(8):2236–2250.

Gokey JJ, Snowball J, Sridharan A, Speth JP, Black KE, Hariri LP, Perl AT, Xu Y, Whitsett JA. (2018) MEG3 is increased in idiopathic pulmonary fibrosis and regulates epithelial cell differentiation. JCI Insight. 3(17):e122490.

Haga CL, Phinney DG (2012). MicroRNAs in the imprinted DLK1-DIO3 region repress the epithelial-to-mesenchymal transition by targeting the TWIST1 protein signaling network. J Biol Chem. 287(51):42695–707.

Hajra KM, Chen DY, Fearon ER. (2002) The SLUG zinc-finger protein represses E-cadherin in breast cancer. Cancer Res. 62(6):1613–8.

Huang HT, Brand OM, Mathew M, Ignatiou C, Ewen EP, McCalmon SA, Naya FJ. (2006) Myomaxin is a novel transcriptional target of MEF2A that encodes a Xin-related alpha-actinin-interacting protein. J Biol Chem. 281(51):39370–9.

Ioannides Y, Lokulo-Sodipe K, Mackay DJ, Davies JH, Temple IK. (2014) Temple syndrome: improving the recognition of an underdiagnosed chromosome 14 imprinting disorder: an analysis of 51 published cases. J Med Genet. 51(8):495–501.

Ito T, Ogawa R, Uezumi A, Ohtani T, Watanabe Y, Tsujikawa K, Miyagoe-Suzuki Y, Takeda S, Yamamoto H, Fukada S. (2013) Imatinib attenuates severe mouse dystrophy and inhibits proliferation and fibrosis-marker expression in muscle mesenchymal progenitors. Neuromuscul Disord. 23(4):349–56.

Joe AW, Yi L, Natarajan A, Le Grand F, So L, Wang J, Rudnicki MA, Rossi FM. (2010) Muscle injury activates resident fibro/adipogenic progenitors that facilitate myogenesis. Nat Cell Biol. 12(2):153–163.

Juan AH, Derfoul A, Feng X, Ryall JG, Dell’Orso S, Pasut A, Zare H, Simone JM, Rudnicki MA, Sartorelli V. (2011) Polycomb EZH2 controls self-renewal and safeguards the transcriptional identity of skeletal muscle stem cells. Genes Dev. 25(8):789–94.

Kameswaran V, Bramswig NC, McKenna LB, Penn M, Schug J, Hand NJ, Chen Y, Choi I, Vourekas A, Won KJ, Liu C, Vivek K, Naji A, Friedman JR, Kaestner KH. (2014) Epigenetic regulation of the DLK1-MEG3 microRNA cluster in human type 2 diabetic islets. Cell Metab. 19(1):135–45.

Kaneko S, Bonasio R, Saldaña-Meyer R, Yoshida T, Son J, Nishino K, Umezawa A, Reinberg D (2014). Interactions between JARID2 and noncoding RNAs regulate PRC2 recruitment to chromatin. Mol Cell. 53(2):290–300.

Kanisicak O, Mendez JJ, Yamamoto S, Yamamoto M, Goldhamer DJ. (2009) Progenitors of skeletal muscle satellite cells express the muscle determination gene, MyoD. Dev Biol. 332(1):131–41.

Kollias HD, McDermott JC. (2008) Transforming growth factor-beta and myostatin signaling in skeletal muscle. J Appl Physiol. 104(3):579–87.

Krauss RS. (2010) Regulation of promyogenic signal transduction by cell-cell contact and adhesion. Exp Cell Res. 316(18):3042–9.

Labialle S, Marty V, Bortolin-Cavaillé ML, Hoareau-Osman M, Pradère JP, Valet P, Martin PG, Cavaillé J. (2014) The miR-379/miR-410 cluster at the imprinted Dlk1-Dio3 domain controls neonatal metabolic adaptation. EMBO J. 33(19):2216–30.

Lamouille S, Xu J, Derynck R. (2014) Molecular mechanisms of epithelial-mesenchymal transition. Nat Rev Mol Cell Biol. 15(3):178–96.

Li Z, Jin C, Chen S, Zheng Y, Huang Y, Jia L, Ge W, Zhou Y. Long non-coding RNA (2017) MEG3 inhibits adipogenesis and promotes osteogenesis of human adipose-derived mesenchymal stem cells via miR-140-5p. Mol Cell Biochem. 433(1-2):51–60.

Liu L, Luo GZ, Yang W, Zhao X, Zheng Q, Lv Z, Li W, Wu HJ, Wang L, Wang XJ, Zhou Q. (2010) Activation of the imprinted Dlk1-Dio3 region correlates with pluripotency levels of mouse stem cells. J Biol Chem. 285(25):19483–90.

Liu L, Cheung TH, Charville GW, Hurgo BM, Leavitt T, Shih J, Brunet A, Rando TA. (2013) Chromatin modifications as determinants of muscle stem cell quiescence and chronological aging. Cell Rep. 4(1):189–204.

Lovett F, Gonzalez I, Salih D, Cobb L, Tripathi G, Cosgrove R, Murrell A, Kilshaw J, Pell J. (2006) Convergence of Igf2 expression and adhesion signalling via RhoA and p38 MAPK enhances myogenic differentiation. J Cell Science. 119:4828–40.

Lu M, Krauss R. (2010) Abl promotes cadherin-dependent adhesion and signaling in myoblasts. Cell Cycle. 9(14): 2737–41.

Lukjanenko L, Karaz S, Stuelsatz P, Gurriaran-Rodriguez U, Michaud J, Dammone G, Sizzano F, Mashinchian O, Ancel S, Migliavacca E, Liot S, Jacot G, Metairon S, Raymond F, Descombes P, Palini A, Chazaud B, Rudnicki MA, Bentzinger CF, Feige JN. (2019) Aging Disrupts Muscle Stem Cell Function by Impairing Matricellular WISP1 Secretion from Fibro-Adipogenic Progenitors. Cell Stem Cell. 24(3):433–446.

Luo Z, Lin C, Woodfin AR, Bartom ET, Gao X, Smith ER, Shilatifard A. (2016) Regulation of the imprinted Dlk1-Dio3 locus by allele-specific enhancer activity. Genes Dev. 30(1):92–101.

Malecova B, Gatto S, Etxaniz U, Passafaro M, Cortez A, Nicoletti C, Giordani L, Torcinaro A, De Bardi M, Bicciato S, De Santa F, Madaro L, Puri PL. (2018) Dynamics of cellular states of fibro-adipogenic progenitors during myogenesis and muscular dystrophy. Nat Commun. 9(1):3670.

Mikovic J, Sadler K, Butchart L, Voisin S, Gerlinger-Romero F, Della Gatta P, Grounds MD, Lamon S. 2018. MicroRNA and Long Non-coding RNA Regulation in Skeletal Muscle From Growth to Old Age Shows Striking Dysregulation of the Callipyge Locus. Front Genet. 9:548.

Mondal T, Subhash S, Vaid R, Enroth S, Uday S, Reinius B, Mitra S, Mohammed A, James AR, Hoberg E, Moustakas A, Gyllensten U, Jones SJ, Gustafsson CM, Sims AH, Westerlund F, Gorab E, Kanduri C (2015). MEG3 long noncoding RNA regulates the TGF-β pathway genes through formation of RNA-DNA triplex structures. Nat Commun. 6:7743.

Nishiyama T, Kii I, Kudo A. (2004) Inactivation of Rho/ROCK signaling is crucial for the nuclear accumulation of FKHR and myoblast fusion. J Biol Chem. 279(45):47311–9.

Ogata T, Kagami M, Ferguson-Smith AC. (2008) Molecular mechanisms regulating phenotypic outcome in paternal and maternal uniparental disomy for chromosome 14. Epigenetics. 3(4):181–7.

Qian P, He XC, Paulson A, Li Z, Tao F, Perry JM, Guo F, Zhao M, Zhi L, Venkatraman A, Haug JS, Parmely T, Li H, Dobrowsky RT, Ding WX, Kono T, Ferguson-Smith AC, Li L. (2016) The Dlk1-Gtl2 Locus Preserves LT-HSC Function by Inhibiting the PI3K-mTOR Pathway to Restrict Mitochondrial Metabolism. Cell Stem Cell. 18(2):214–28.

Rathbone CR, Yamanouchi K, Chen XK, Nevoret-Bell CJ, Rhoads RP, Allen RE. (2011) Effects of transforming growth factor-beta (TGF-β1) on satellite cell activation and survival during oxidative stress. J Muscle Res Cell Motil. 32(2):99–109.

Sartori R, Schirwis E, Blaauw B, Bortolanza S, Zhao J, Enzo E, Stantzou A, Mouisel E, Toniolo L, Ferry A, Stricker S, Goldberg AL, Dupont S, Piccolo S, Amthor H, Sandri M. (2013) BMP signaling controls muscle mass. Nat Genet. 45(11):1309–18.

Schaum, N., Karkanias, J., Neff, N.F. et al. 2018. Single-cell transcriptomics of 20 mouse organs creates a *Tabula Muris*. Nature 562,367–372.

Segalés J, Perdiguero E, Muñoz-Cánoves P. (2016) Regulation of Muscle Stem Cell Functions: A Focus on the p38 MAPK Signaling Pathway. Front Cell Dev Biol. 4:91.

Shin YJ, Kwon ES, Lee SM, Kim SK, Min KW, Lim JY, Lee B, Kang JS, Kwak JY, Son YH, Choi JY, Yang YR, Kim S, Kim YS, Jang HC, Suh Y, Yoon JH, Lee KP, Kwon KS. (2020) A subset of microRNAs in the Dlk1-Dio3 cluster regulates age-associated muscle atrophy by targeting Atrogin-1. J Cachexia Sarcopenia Muscle. doi: 10.1002/jcsm.12578.

Snyder CM, Rice A, Estrella N, Held A, Kandarian SC, Naya FJ (2013). MEF2A regulates the Gtl2-Dio3 microRNA mega-cluster to modulate WNT signaling in skeletal muscle regeneration. Development. 140(1): p. 31–42.

Soleimani VD, Yin H, Jahani-Asl A, Ming H, Kockx CE, van Ijcken WF, Grosveld F, Rudnicki MA. (2012) Snail regulates MyoD binding-site occupancy to direct enhancer switching and differentiation-specific transcription in myogenesis. Mol Cell. 47(3):457–68.

Sousa-Victor P, Gutarra S, García-rat L, Rodriguez-Ubreva J, Ortet L, Ruiz-Bonilla V, Jardí M, Ballestar E, González S, Serrano AL, Perdiguero E, Muñoz-Cánoves P. (2014) Geriatric muscle stem cells switch reversible quiescence into senescence. Nature. 506(7488):316–21.

Stadtfeld M, Apostolou E, Akutsu H, Fukuda A, Follett P, Natesan S, Kono T, Shioda T, Hochedlinger K. (2010) Aberrant silencing of imprinted genes on chromosome 12qF1 in mouse induced pluripotent stem cells. Nature. 465(7295):175–81.

Terashima M, Tange S, Ishimura A, Suzuki T (2017). MEG3 Long Noncoding RNA Contributes to the Epigenetic Regulation of Epithelial-Mesenchymal Transition in Lung Cancer Cell Lines. J Biol Chem. 292(1):82–99.

Tierling S, Dalbert S, Schoppenhorst S, Tsai CE, Oliger S, Ferguson-Smith AC, Paulsen M, Walter J. (2006) High-resolution map and imprinting analysis of the Gtl2-Dnchc1 domain on mouse chromosome 12. Genomics. 87(2):225–35.

Uezumi A, Ikemoto-Uezumi M, Tsuchida K (2014). Roles of nonmyogenic mesenchymal progenitors in pathogenesis and regeneration of skeletal muscle. Front Physiol. 5:68.

Ungefroren H, Witte D, Lehnert H. (2018) The role of small GTPases of the Rho/Rac family in TGF-β-induced EMT and cell motility in cancer. Dev Dyn. 247(3):451–461.

Woodhouse S, Pugazhendhi D, Brien P, Pell JM. (2013) Ezh2 maintains a key phase of muscle satellite cell expansion but does not regulate terminal differentiation. J Cell Sci. 126(Pt 2):565–79.

Wosczyna MN, Rando TA. (2018) A Muscle Stem Cell Support Group: Coordinated Cellular Responses in Muscle Regeneration. Dev Cell. 46(2):135–143.

Wosczyna MN, Konishi CT, Perez Carbajal EE, Wang TT, Walsh RA, Gan Q, Wagner MW, Rando TA. (2019) Mesenchymal Stromal Cells Are Required for Regeneration and Homeostatic Maintenance of Skeletal Muscle. Cell Rep. 27(7):2029–2035.

Wüst S, Dröse S, Heidler J, Wittig I, Klockner I, Franko A, Bonke E, Günther S, Gärtner U, Boettger T, Braun T (2018) Metabolic Maturation during Muscle Stem Cell Differentiation Is Achieved by miR-1/133a-Mediated Inhibition of the Dlk1-Dio3 Mega Gene Cluster. Cell Metab. 27(5):1026–1039.

Yen, Y-P, Hsieh, W-F, Tsai, Y-Y, Lu, Y-L, Liau, E.S., Hsu, H-C, Chen Y-C, Liu, T-C, Chang, M, Li, J, Lin, S-P, Hung, J-H, Chen, J-A. (2018) Dlk1-Dio3 locus-derived lncRNAs perpetuate postmitotic motor neuron cell fate and subtype identity. eLife 7, e38080.

Yu F, Geng W, Dong P, Huang Z, Zheng J. (2018). LncRNA-MEG3 inhibits activation of hepatic stellate cells through SMO protein and miR-212. Cell Death Dis. 9(10):1014.

Yusuf F, Brand-Saberi B. (2006) The eventful somite: patterning, fate determination and cell division in the somite. Anat Embryol (Berl). 211 Suppl 1:21–30.

Zhao J, Ohsumi TK, Kung JT, Ogawa Y, Grau DJ, Sarma K, Song JJ, Kingston RE, Borowsky M, Lee JT. (2010) Genome-wide identification of polycomb-associated RNAs by RIP-seq. Mol Cell. 40(6):939–53.

Zhou Y, Zhong Y, Wang Y, Zhang X, Batista DL, Gejman R, Ansell PJ, Zhao J, Weng C, Klibanski A. (2007) Activation of p53 by MEG3 non-coding RNA. J Biol Chem. 282(34):24731–42.

Zhou, Y., Cheunsuchon, P., Nakayama, Y., Lawlor, M.W., Zhong, Y., Rice, K.A., Zhang, L., Zhang, X., Gordon, F.E., Lidlov, H.G.W., et al. (2010). Activation of paternally expressed genes and perinatal death caused by deletion of the *Gtl2* gene. Development 137, 2643–2652.

